# RNA sequencing depth guidelines for the study of alternative splicing

**DOI:** 10.1101/2024.10.09.617406

**Authors:** Olga Tsoy, Sabine Ameling, Sören Franzenburg, Markus Hoffmann, Lina Liv-Willuth, Hye Kyung Lee, Ludwig Knabl, Priscilla A. Furth, Uwe Völker, Lothar Hennighausen, Jan Baumbach, Tim Kacprowski, Markus List

**Author notes:** Joint Last Authors (Tim Kacprowski, Markus List).

## Abstract

A key parameter in the experimental design of RNA-seq projects is the choice of sequencing depth. Considering a limited budget, one needs to find a tradeoff between the number of samples and the sensitivity of the analysis, particularly concerning lowly expressed genes. While previous studies have proposed a lower bound for the comprehensive analysis of differential gene expression, for the analysis of alternative splicing, it has only been proposed for human adipose tissue. However, alternative splicing differs across tissues and conditions. We analyzed publicly available and newly generated deep-sequenced paired-end RNA-seq samples (between 150 and >500 million reads, read length 50-150 bp) from human buffy coat cells and diverse sets of tissues, including gluteal subcutaneous fat, heart, and hypothalamus. Our results show that the sequencing depth typically used in published cohorts is not sufficient to comprehensively capture the landscape of alternative splicing. This motivates the use of deeper sequencing or long-read technologies in future studies. Toward this goal, we offer guidelines for choosing a suitable sequencing depth.

**GRAPHICAL ABSTRACT:** 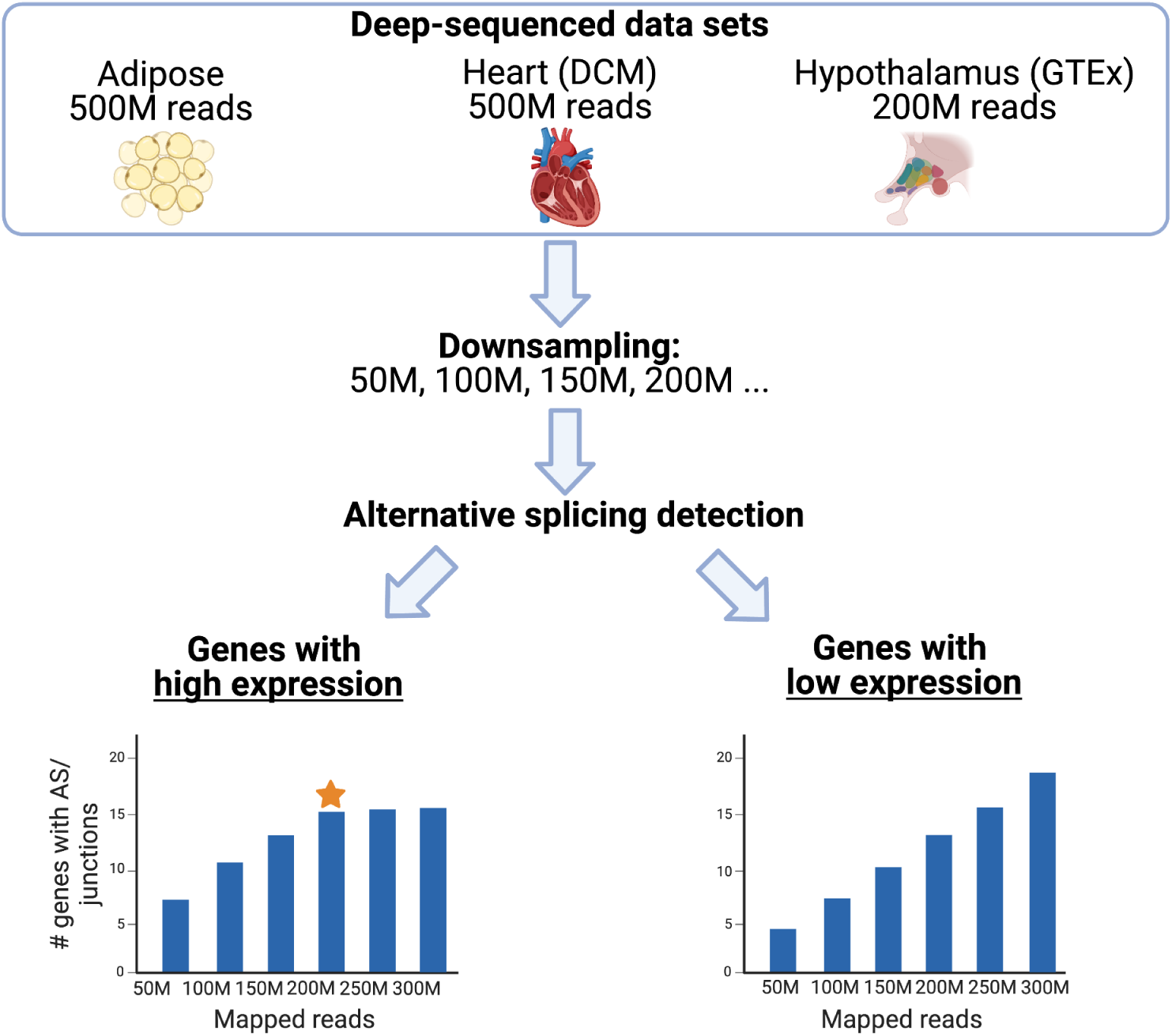

## INTRODUCTION

Short-read RNA sequencing is commonly used to study alternative splicing (AS) due to low sequencing error, the possibility of counting the abundance of sequences, and cost-effectiveness. Robust detection of AS events depends on the sequencing depth, which is especially crucial if a researcher aims to detect low-abundant transcripts (1). Yet, it remains unclear how many reads are required for comprehensive coverage of AS in human samples. Considering the tradeoff between sequencing depth and sample number, guidelines for the appropriate experimental design guidelines are needed.

The impact of sequencing depth on AS detection has been partially tackled in evaluating AS tools. SUPPA2 evaluation demonstrated that at 120M reads, the proportion of true positive detections reaches almost 100% for exon skipping but 90% for alternative splice site events (2). In the Cufflinks paper, the authors demonstrated that the recall suffered for low abundant transcripts even at 100M reads (3). When evaluating CASH (4), the authors demonstrated a notable increase in the performance of AS detection tools with the increase of sequencing depth from 50M to 100M reads. We also partially addressed this question while evaluating the AS simulation tool ASimulatoR (5). In the study, we simulated the data set with 200M reads. We observed the increase in AS detection performance between 50M and 100M reads, as well as a steadily increasing number of detected AS events with a sequencing depth of more than 100M reads. In our benchmark of AS detection tools (6), we observed that most AS detection tools demonstrate high precision regardless of the sequencing depth but recall further increases even at 200M reads.

However, all of the studies mentioned above use simulated data, where the number of AS events is chosen artificially and might not reach the complexity of the biological data. Moreover, most of the studies simulated less than 100M reads (Supplementary Table S1). Finally, the primary focus of the AS detection tools evaluations is to demonstrate the performance and robustness of a tool. Thus, these studies do not answer the question of the RNA sequencing depth threshold for comprehensive AS detection.

Several studies addressed the sequencing depth threshold question for gene expression analysis. Estimates for the appropriate sequencing depth for differential gene expression analysis range from 200M (7) to 300M reads (8) for the human genome. Bass *et al.* stated that 70% of all studies are “undersequenced” for thorough differential gene expression analysis (9). The number of studies concerning AS is still limited. The SEQC Consortium investigated the discovery of exon-exon junctions and demonstrated that new junctions can still be detected even at 1 billion reads (10). Lihuh et al. investigated the limitation of RNA-Seq for detecting rare transcripts using the data from *D. melanogaster* with spike-ins (11). They estimated that at least 68M uniquely aligned single-end 36-bp reads are needed to detect transcripts present at one copy per cell. Considering the larger size of the human genome and the larger number of genes, this number should be much larger for human data analysis. Hardwick et al. (12) presented an interesting approach to adding the synthetic transcriptome with known transcript sequences (sequencing spike-ins, ‘sequins’). They observed that even at 100M reads, they could not detect the complete assembly. Finally, one study investigated this issue in human adipose tissue and suggested a lower bound of 150M paired reads (8). However, considering the tissue-specificity of AS, an in-depth analysis of the necessary sequencing depth for its robust detection in different tissues is missing.

We analyzed RNA-seq data from diverse human tissues and conditions, considering published sources as well as very deeply sequenced heart tissue samples (> 500M reads). Since the sensitivity of AS detection depends on the expression level of a gene, we considered genes with low expression levels (TPM < 0.1, 0.1 ≤ TPM < 0.5, and 0.5 ≤ TPM < 1) to focus on low abundant transcripts, and high expression levels (1 ≤ TPM < 10, TPM ≥ 10) to suggest different lower bounds depending on a gene expression level.

## MATERIAL AND METHODS

The poly(A) RNA-seq datasets used in the study are listed in Table 1. RNA-seq data of human buffy coat cells isolated from whole blood from patients with SARS-CoV-2 variant Alpha (GSE190680 (13)) were obtained from Gene Expression Omnibus (GEO) (14). This cohort contains RNA-seq samples sequenced at various sequencing depths (from 50M to 150M) and collected at different time points after the first symptoms (13): 1-5 days, 10-14 days, and 15-30 days. RNA from the samples of the SARS-CoV-2 data had a quality RNA integrity (RIN) score between 8.8 and 10.0 with a mean of 9.82. The RIN value was measured by Agilent Bioanalyzer 2100.

**Table 1.** RNA-seq data sets used in the study.

| Data set | Source | #of samples | #of paired reads (in M) | Read length (bp) |
| --- | --- | --- | --- | --- |
| Human buffy coat white blood cells from SARS-CoV-2 infected patients | GSE190680 | 97 | 60 - 150 | 150 |
| Adipose | GSE46323 | 2 | 455-519 | 101 |
| Heart (DCM) | This study<sup>1</sup> | 4 | 519 - 527 | 101 |
| Hypothalamus | GTEX (V8) | 102 | 10 - 236 | 76 |
| Breast cancer (primary tissue) | TCGA | 100 | 82-189 | 50 |

<sup>1</sup>Due to the restrictions of the patient consent, data can only be available upon request from University Medicine Greifswald

Total RNA was extracted from human frozen endomyocardial biopsies using TRIzol (ThermoFisher Scientific, Waltham, MA, USA), followed by DNA digestion purification steps using the RNAase-Free DNase I Kit and RNA Clean-Up and Concentration kit according to the manufacturer’s instructions (both Norgen Biotek Corp., Thorold, ON, Canada). The quality of the extracted RNA was assessed using the Tape Station 4200 (Agilent, Santa Clara, CA, USA). All four samples exhibited a mean RIN of 8.5. Subsequently, a PolyA library was prepared based on 500 ng total RNA per sample using the TruSeq stranded mRNA kit (Illumina, San Diego, CA, USA). For sequencing, 200 pM of the final library was loaded on the NovaSeq 6000 System, S1 FlowCell (2 x 100 bp paired-end reads). A sequencing depth of approximately 500 Mio clusters per sample was obtained and recorded at the Competence Centre for Genomic Analysis in Kiel.

RNA-seq samples from human adipose tissue before and after treatment (exposure to endotoxin) (GSE46323) were also downloaded from GEO. RNA-seq data from the hypothalamus (SRR1443713) has been downloaded from the Genotype-Tissue Expression (GTEx) project(15). RNA-Seq data from breast cancer (primary tissue) have been downloaded from the NIH National Cancer Institute GDC data portal https://portal.gdc.cancer.gov/projects/TCGA-BRCA (sample-IDs: https://doi.org/10.6084/m9.figshare.22361005.v1).

All RNA-seq data (except the data from TCGA) were obtained as raw reads and aligned to the human reference genome GRCh38.p13 with STAR 3.7 (default parameters, 2-pass mode) (16) as this tool has been demonstrated to be the optimal tool for a human genome analysis (6, 17). The TCGA data has been obtained as alignments in the bam format. The gene counts were calculated by featureCounts from Subread 2.0 (18) and normalized to transcripts per million (TPM). Downsampling of aligned reads was performed with samtools 1.7 (19) using the subsampling option (“view -s”). The subsampling procedure in samtools randomly eliminates RNA-Seq reads down to the requested amount. By design, the downsampling procedure of samtools takes into account only mapped reads. Thus, for all analyses in this study, we used the number of mapped reads as an indication of sequencing depth. The number of mapped reads of the analyzed samples has been obtained from the STAR log files or samtools flagstat output for the TCGA samples. The number of expressed junctions has been obtained using regtools 0.5.2 (20).

Among the published tools that can detect AS events in one condition, we chose MAJIQ (21), which showed high precision, acceptable recall, and good run time in a recently published benchmark (6). MAJIQ also detects both known and novel AS events. We ran an AS analysis using MAJIQ 2.1 with default parameters. While MAJIQ 2.1 uses the notation of “local splicing variant” - alternative junctions bound to the same exon - and not AS event types (e.g., exon skipping, intron retention, etc), we did not count AS events but genes with AS and junctions involved in AS. Also, we considered only those ‘local splicing variants’ with the percent splice-in (PSI) values more than 0.05 and less than 0.95 to decrease the number of false-positive AS events detections and focus on biologically relevant ones. The number of genes with AS and the number of junctions involved in AS were calculated and visualized by in-house Python scripts.

The Mann-Kendall Trend test has been performed using the pymannkendall 1.4.3 Python package (https://pypi.org/project/pymannkendall/). The reported Mann-Kendall Trend slope presents a magnitude of the monotonic trend. The GTEx Hypothalamus dataset lacks enough data points to run the trend test.

The functional enrichment for the genes with AS detected only at 200M reads has been performed using g:Profiler (22). The AS-aware enrichment was performed using NEASE (23) using the list of exon skipping events as an input. The events are visualized via the sashimi tool (24).

The cost for detecting an additional gene with AS/alternative junction with an increasing number of mapped reads has been calculated according to the prices from the Competence Centre for Genome Analysis Kiel (CCGA Kiel, https://ccga.uni-kiel.de). We divided the cost of sequencing for each additional 50M reads by the number of genes with AS/junctions identified with this new portion of reads.

All Python scripts used to generate the figures as well as the MAJIQ raw results and summary files are available at https://github.com/OlgaVT/RNA-Seq_Depth_for_AS and https://doi.org/10.5281/zenodo.11655945.

## RESULTS

### SARS-CoV-2 cohort

We investigated how the number of detected genes with AS events/junctions involved in AS events depends on a sequencing depth in real-world data. However, a sequencing depth higher than 50-100 M reads is uncommon for large RNA-Seq datasets. The SARS-CoV-2 cohort used in the study was chosen because of the deeper than usual amount of reads (up to 150M) (13).

For each sample from the SARS-CoV-2 cohort, we calculated how many genes with AS events and how many junctions involved in AS events were detected. We considered genes with low expression (TPM < 0.1, 0.1 ≤ TPM < 0.5, and 0.5 ≤ TPM < 1) and high expression (1 ≤ TPM < 10, TPM ≥ 10) and AS events with 0.05 < PSI < 0.95. We examined how the number of detected genes/junctions depends on the number of uniquely mapped reads reported by STAR (Figure 1). The total number of genes and junctions in each category is in Supplementary Table S2. We tested if there was an increasing trend using the Mann-Kendall Trend test and calculated the increase in the mean number of detections (Table 2) Percentage of new detections was calculated as the ratio between the mean value of detections at a certain sequencing depth and the mean total value of genes/junctions per expression category.

**Figure 1:**
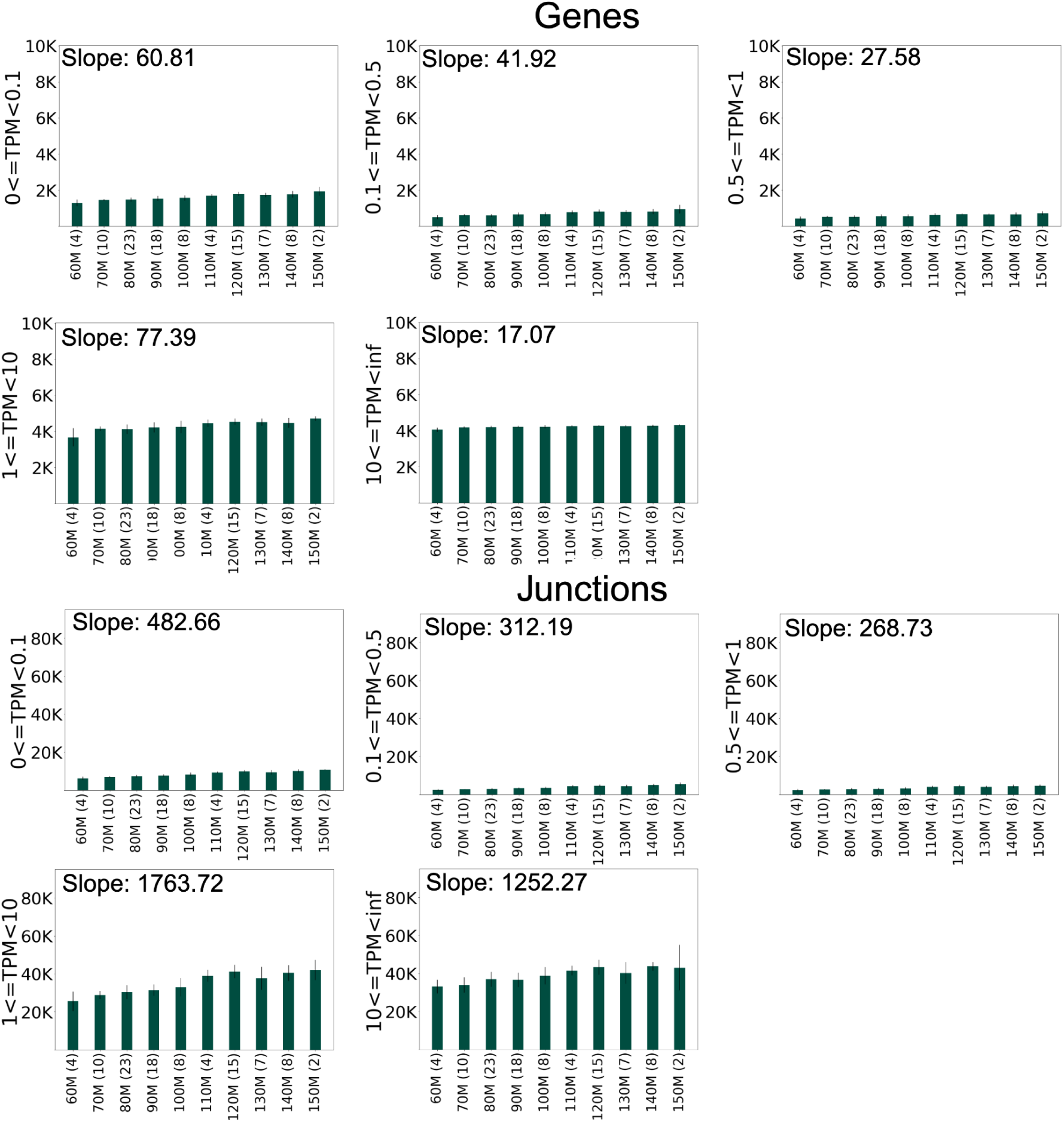
The number of genes with AS and junctions involved in AS detected from SARS-CoV-2 RNA-seq data with various numbers of mapped reads. The bar plots demonstrate the mean number of genes/junctions involved in AS. Slopes (a magnitude of the monotonic trend) are indicated for significant (p-value < 0.05) Mann-Kendall Trend Test results.

**Table 2.** The increase of the mean number and the percentage of genes and junctions involved in alternative splicing with increasing sequencing depth in the SARS-Cov2 dataset. The column ‘0-60 M’ indicates the mean number and the percentage of genes/junctions detected in the samples with a sequencing depth of less than 60 M reads. The column ‘150M+’ indicates the mean number and the percentage of genes/junctions detected in the samples with the highest sequencing depth of more than 150M reads. Grey shading indicates an increase of less than 1% new detections.

| TPM | 0-60M | 60-70 | 70-80 | 80-90 | 90-100 | 100-110 | 110-120 | 120-130 | 130-140 | 140-150 | 150M+ | Mean total # of genes/junctions |
| --- | --- | --- | --- | --- | --- | --- | --- | --- | --- | --- | --- | --- |
|  | **Genes** |  |  |  |  |  |  |  |  |  |  |  |
| < 0.1 | 1287.8<br>(20.9%) | +161.3<br>(+2.6%) | +19.9<br>(+0.3%) | +63.4<br>(+1.0%) | +25.4<br>(+0.4%) | +132.6<br>(+2.2%) | +91.2<br>(+1.5%) | -56.7<br>(-0.9%) | +43.9<br>(+0.7%) | +165.2<br>(+2.7%) | 1932.5<br>(31.4%) | 6149 |
| [0.1, 0.5) | 498<br>(11.8%) | +105.4<br>(+2.5%) | -14.7<br>(0.4%) | +54.3<br>(+1.3%) | +19.4<br>(+0.5%) | +107.4<br>(2.5%) | +48.1<br>(+1.1%) | -19.6<br>(-0.5%) | +6.9<br>(+0.2%) | +128.3<br>(+3%) | 933.5<br>(22.2%) | 4213 |
| [0.5, 1) | 416<br>(20.7%) | +104.7<br>(5.2%) | -7.9<br>(-0.4%) | +29.5<br>(+1.5%) | +10.8<br>(+0.5%) | +77.9<br>(+3.9%) | +25.7<br>(+1.3%) | -13.4<br>(-0.7%) | +3.2<br>(+0.2%) | +68.5<br>(+3.4%) | 715<br>(35.6%) | 2006 |
| [1, 10) | 3671.8<br>(50.6%) | +473.4<br>(+6.5%) | -25.5<br>(-0.4%) | +97.5<br>(+1.3%) | +27.3<br>(+0.4%) | +214.1<br>(+3%) | +77.2<br>(+1.1%) | -31.4<br>(-0.4%) | -30.2<br>(-0.4%) | +232.2<br>(+3.2%) | 4706.5<br>(64.9%) | 7254 |
| ≥ 10 | 4048.3<br>(79.2%) | +134.9<br>(+2.6%) | +16.8<br>(+0.3%) | +19.7<br>(+0.4%) | +5.4<br>(+0.1%) | +28.8<br>(+0.6%) | +25.5<br>(+0.5%) | -29.9<br>(-0.6%) | +19.4<br>(+0.4%) | +37.3<br>(+0.7%) | 4306<br>(84.2%) | 5112 |
|  | **Junctions** |  |  |  |  |  |  |  |  |  |  |  |
| < 0.1 | 6283.3<br>(25.9%) | +716.1<br>(+2.9%) | +348.2<br>(+1.4%) | +383.6<br>(+1.6%) | +528.0<br>(+2.2%) | +1016.5<br>(+4.2%) | +673.7<br>(+2.8%) | -521.9<br>(-2.1%) | +686.4<br>(+2.8%) | +717.5<br>(+3.0%) | 10831.5<br>(44.6%) | 24302 |
| [0.1, 0.5) | 2407<br>(7.5%) | +429.4<br>(+1.3%) | +90.3<br>(+0.3%) | +304.4<br>(+0.9%) | +272.1<br>(+0.8%) | +786.5<br>(+2.5%) | +458.5<br>(+1.4%) | -388.8<br>(-1.3%) | +440.0<br>(+1.4%) | +602.7<br>(+1.9%) | 5402.5<br>(16.8%) | 32095 |
| [0.5, 1) | 2212.8<br>(10.1%) | +444.8<br>(+2.0%) | +92.7<br>(+0.4%) | +185.5<br>(+0.9%) | +174.2<br>(+0.8%) | +948.25<br>(+4.3%) | +368.7<br>(+1.7%) | -418.1<br>(-1.9%) | +312.6<br>(+1.4%) | +415.0<br>(+1.9%) | 4736.5<br>(21.7%) | 21823 |
| [1, 10) | 25612.5<br>(17.7%) | +3267.7<br>(+2.3%) | +1491.1<br>(+1.0%) | +1043.3<br>(+0.7%) | +1583.0<br>(+1.1%) | +5902.9<br>(4.1%) | +2393.7<br>(+1.7%) | -3573.6<br>(-2.5%) | +2731.9<br>(+1.9%) | +1544.5<br>(1.1%) | 41997<br>(29.0%) | 144588 |
| $\geq 10$ | 31105.5<br>(21.4%) | 767.3<br>(+0.5%) | +3083.7<br>(+2.0%) | -321.2<br>(-0.2%) | +2191.5<br>(+1.4%) | +2685.5<br>(+1.7%) | +1760.2<br>(+1.1%) | -3023.5<br>(-2.0%) | +3586.8<br>(+2.3%) | -830.3<br>(-0.5%) | 43005.5<br>(27.8%) | 154833 |

The Mann-Kendall Test confirmed the increasing trend for all expression levels - all calculated slopes are positive (Figure 1). Table 2 shows that for almost all gene expression categories (except TPM > 10), the increase from 60M to 150M reads results in about 10-15% new detections. Even an increase from 140M to 150M still results in more than 1% of new detections.

Thus, to detect AS events with 0.05 < PSI < 0.95 in the genes with TPM < 10, a sequencing depth of 150M reads is insufficient. However, for very highly expressed genes (TPM > 10), the threshold might be around 70M reads.

The trend is similar for the number of detected junctions involved in AS (Table 2). For junctions in genes with TPM < 10, the increase from 60-150M results in about 12-19% new detections. Also, the increase from 140M to 150M reads still results in more than 1% new detections, confirming that 150M reads are insufficient for covering genes with low expression in AS analysis. However, for junctions in very highly expressed genes, the increase declines around 110-120M reads, indicating a much lower threshold for this gene expression category.

While we observe a stable positive trend, the values for the number of genes/junctions involved in AS events show considerable variation across samples. This motivated us to visually investigate and confirm for several genes that the RNA-Seq data does support the expression of junctions that could be detected only in deeply sequenced samples and that the corresponding events do not merely represent false positive AS event detections (Supplementary Figure S1).

### Deep-sequenced cohorts

After the analysis of the SARS-CoV-2 cohort, the question remains: how will the number of AS event detections increase with the sequencing depth of more than 150M reads? We searched for datasets with more than 150M reads and discovered two published datasets with paired-end mRNA RNA-Seq samples: two samples from the publicly available RNA-seq dataset from the human adipose tissue and one hypothalamus RNA-seq sample from the GTEx cohort. We additionally deeply sequenced four samples from the heart tissues of patients suffering from dilated cardiomyopathy (DCM).

We examined how the robustness and sensitivity of AS detection change with an increasing number of mapped reads (Figure 2). We downsampled each sample in the Adipose and Heart (DCM) data sets 100 times to 300, 250, 200, 150, 100, and 50M reads (Figure 3), and Hypothalamus dataset to 200, 150, 100, and 50M (Supplementary Figure S2). As before, we divided the detected genes into groups according to the expression level: low expression levels (TPM < 0.1, 0.1 ≤ TPM < 0.5, and 0.5 ≤ TPM < 1), and high expression levels (1 ≤ TPM < 10, TPM ≥ 10). Then we counted the number of detected genes/junctions involved in AS with 0.05 < PSI < 0.95. The total number of genes in each category is in Supplementary Table S2. In line with the SARS CoV-2 data set analysis, we applied the Mann-Kendall Trend test to detect the significance of the trend and the sign of a slope and calculated the increase in the number of detections (Table 3, Supplementary Tables S4-S6).

**Figure 2:**
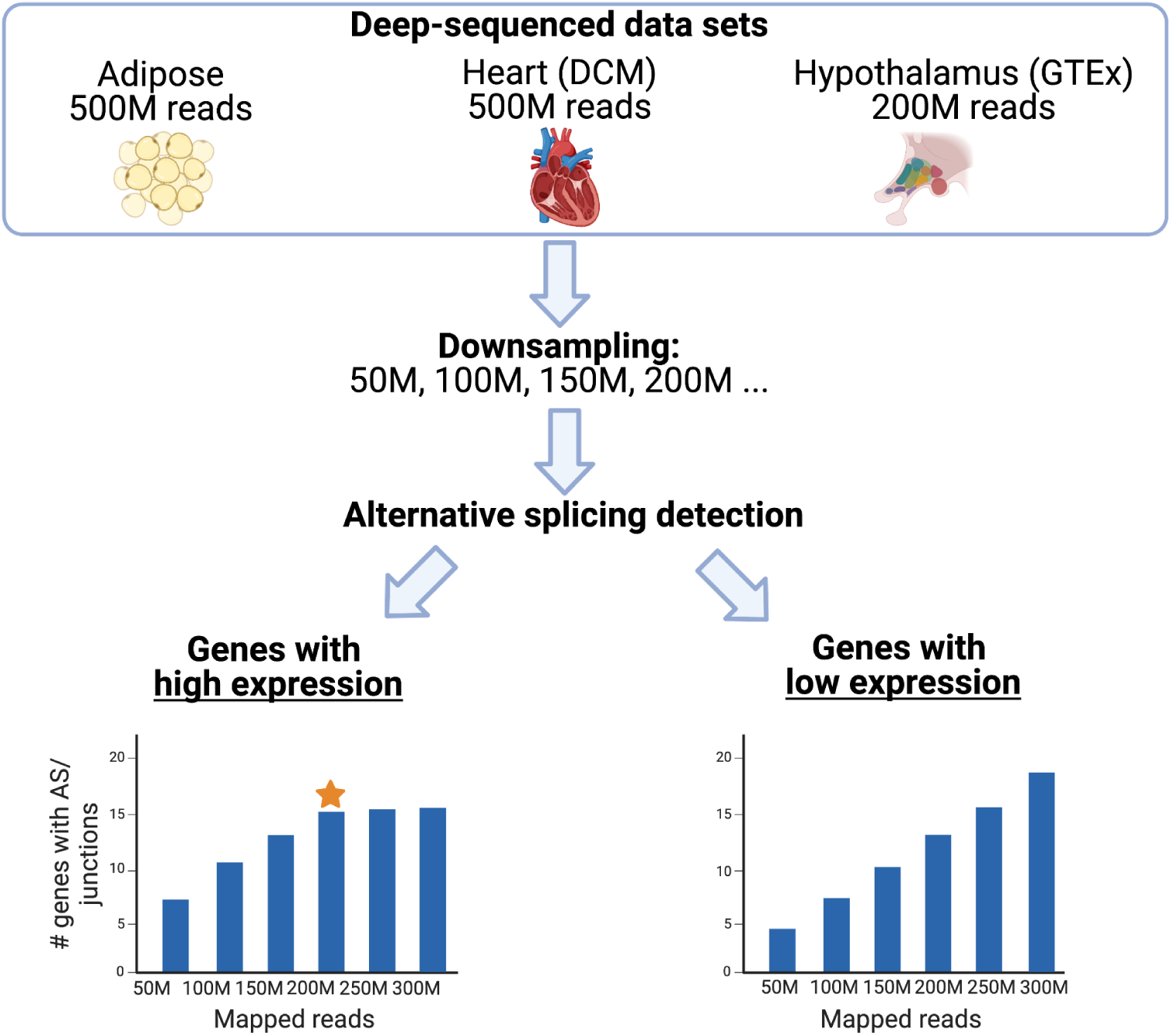
The strategy to estimate the number of mapped reads needed for AS analysis from three deep-sequenced data sets. We expect that new AS events will be steadily detected for genes with low expression due to an increase in sensitivity. In contrast, for highly expressed genes a saturation effect is expected. The orange star marks the threshold for the sequencing depth we aim to identify. Note that the bars shown here are only meant for illustration and do not represent the results. Created with BioRender.

**Figure 3:**
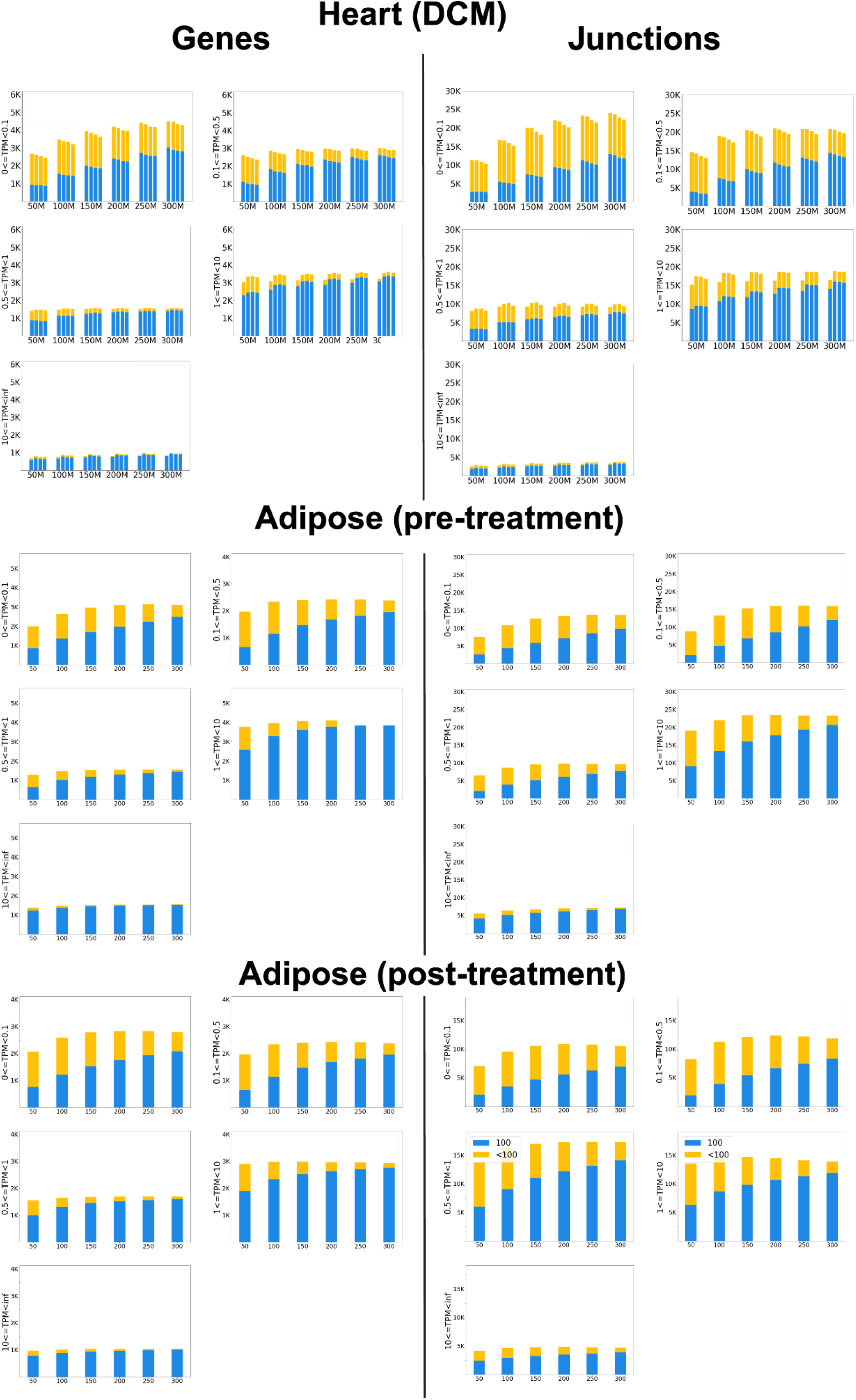
The number of genes and junctions involved in AS detected in 100 (blue), and less than 100 (yellow) out of 100 subsamples Together blue and yellow bars represent the number of genes and junctions involved in AS detected in at least one subsample. Each stacked bar depicts one sample. The slopes for significant (p-value < 0.05) results of the Mann-Kendall Trend Test are in Supplementary Table S3.

**Table 3.**
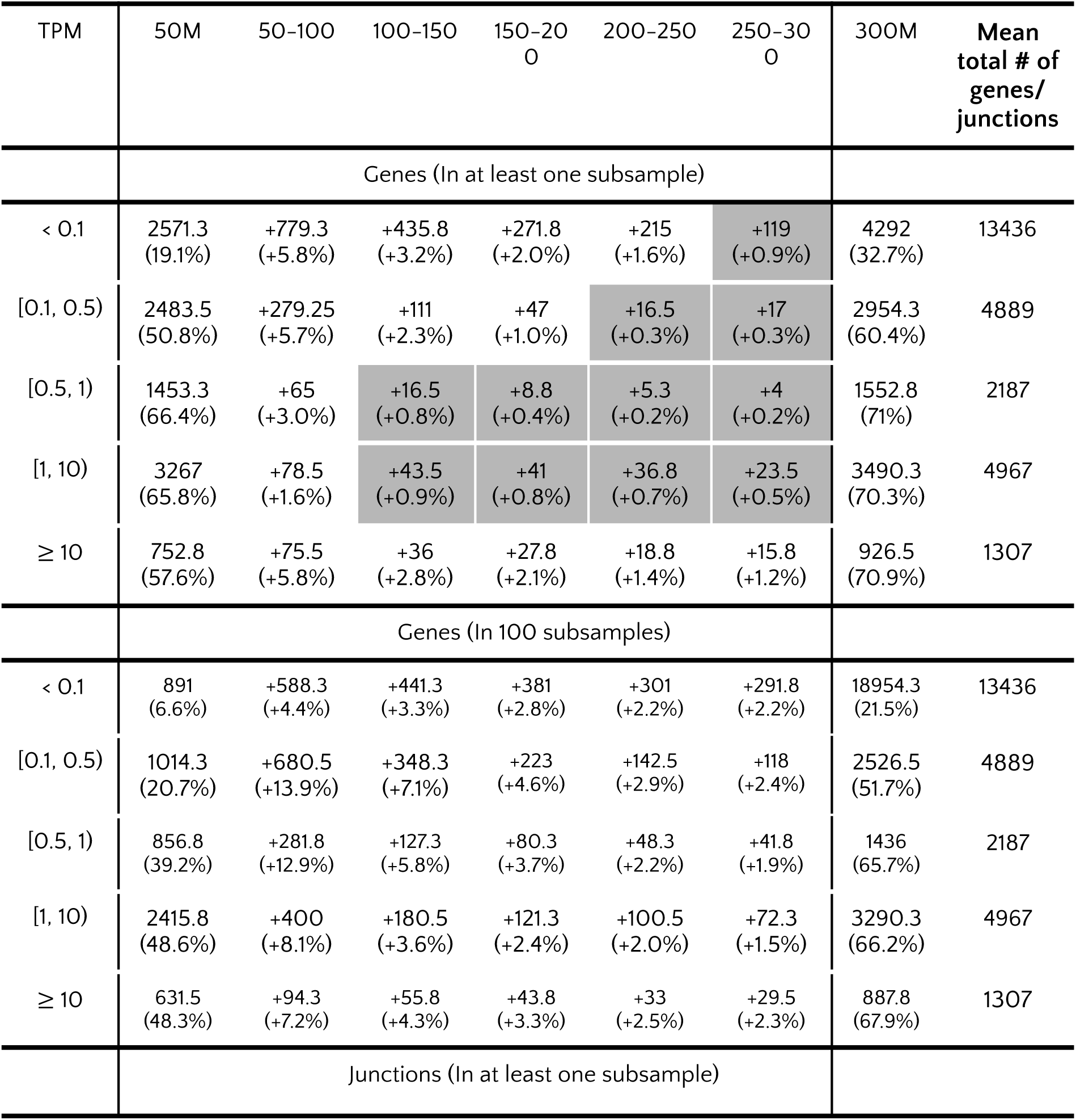
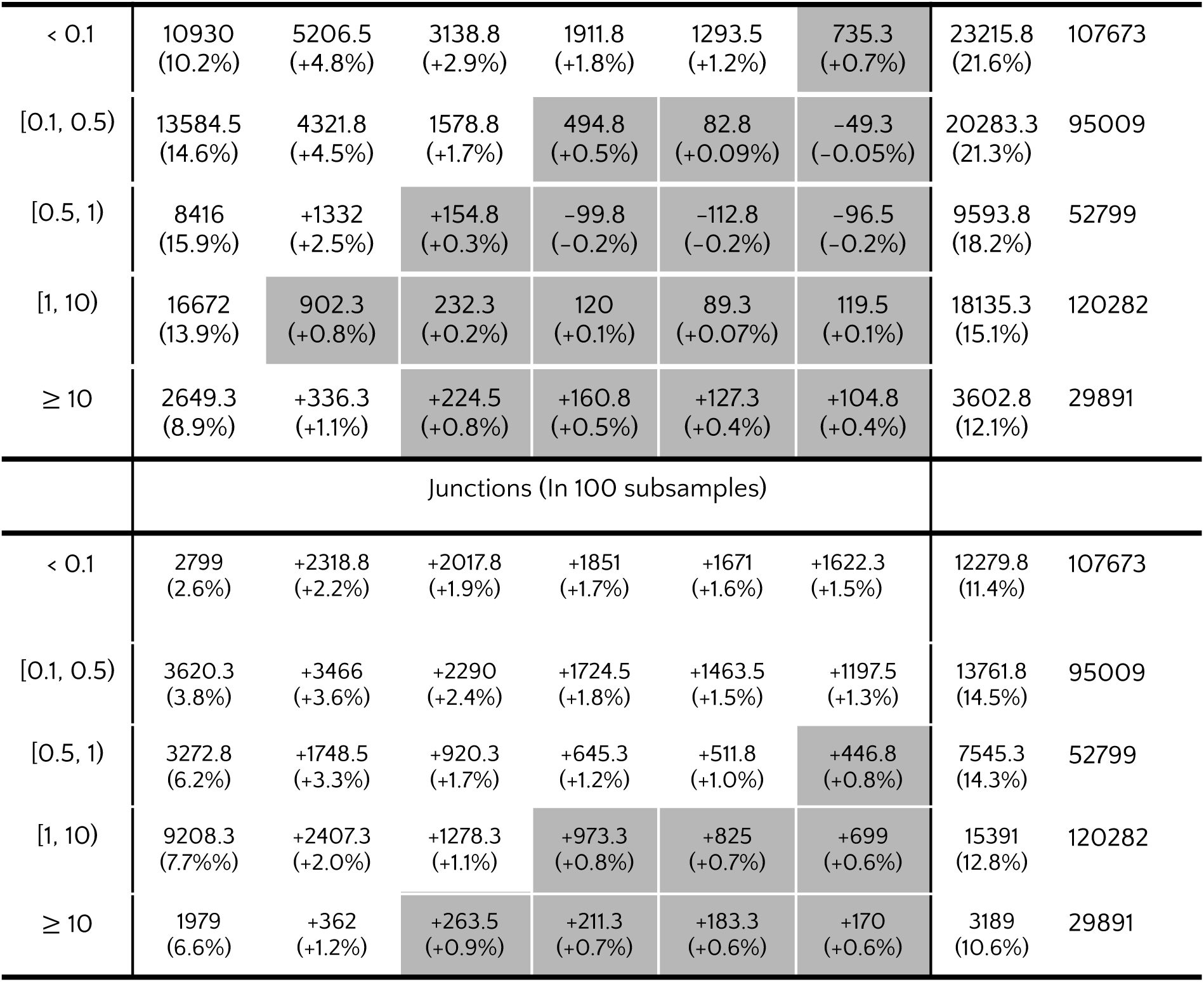
The increase of the mean number and the percentage of genes and junctions involved in AS events with increasing sequencing depth in the Heart (DCM) dataset. Column ‘50 M’ indicates the mean number and the percentage of genes/junctions detected in the samples with a sequencing depth of less than 50 M reads. The column ‘300M’ indicates the mean number and the percentage of genes/junctions detected in the samples with the highest sequencing depth of 300M reads. Grey cells indicate an increase of less than 1%.

We observed differences in how the number of genes with AS/junctions involved in AS changes with an increasing sequencing depth between genes/junctions. Specifically, we compared AS changes consistently detected in all 100 subsamples (Figure 3, blue bar plots) to changes observed in at least one subsample (Figure 3, blue + yellow bar plots). For genes/junctions detected consistently in all 100 subsamples, the Mann-Kendall Trend test reported a significantly increasing trend for all datasets (Supplementary Table S3, “in 100 subsamples”). However, for genes/junctions detected in at least one subsample (Supplementary Table S3, “in at least one subsample”), the trend is not significant in the Adipose (pre-treatment) data set for lowly expressed (0.5 ≤ TPM < 1) genes and for junctions from lowly expressed (0.5 ≤ TPM < 1) and highly expressed (1 ≤ TPM < 10) genes. The trend is also not significant in the Adipose (post-treatment) data set for lowly expressed (TPM < 0.1, 0.1 ≤ TPM < 0.5, and 0.5 ≤ TPM < 1) and highly expressed (1 ≤ TPM < 10) genes with AS and for junctions involved in AS from all gene expression levels except 0.5 ≤ TPM < 1.

Next, for most datasets, for genes with AS/junctions involved in AS found in 100 subsamples (Table 3, Supplementary Tables S4-6, “In 100 subsamples”), we observe more than 1 % of new detections of genes/junctions even with the increase of sequencing depth from 250M to 300M reads. The only exception is the number of junctions involved in AS from the Heart (DCM) dataset (Table 3, “Junctions (In 100 subsamples)). Thus, with higher sequencing depth, more AS events can be robustly found in the data set. However, to confirm all AS events, a sequencing depth higher than 300M is needed.

On the other hand, the increase of genes with AS/junctions involved in AS for those found in at least one subsample slows down - above a certain sequencing depth, there are less than 1% of new detections (Table 3, Tables S4-6, “In at least one subsample”, grey shading). In this case, we suggest the threshold of 150-200M reads for genes/junctions in the low expression category (TPM < 0.1, 0.1 ≤ TPM < 0.5, and 0.5 ≤ TPM < 1) and of 100-150M reads for genes/junctions in the high expression category (1 ≤ TPM < 10, TPM ≥ 10).

### The biological relevance of AS events found at 200M reads

We investigated the biological relevance of the AS events found only at 200M reads using various approaches. First, we extracted genes with AS detected in all 100 samples exclusively at 200M reads but not in 50M, 100M, and 150M, and performed functional gene enrichment with g:Profiler (22). AS events unique to 200M reads in highly expressed genes (TPM > 1) in the adipose sample after endotoxin treatment are enriched in the category ‘response to organic substance’ (Supplementary Figure S3), indicating that higher sequencing depth might reveal additional information about the response mechanism to endotoxin.

One of the datasets presents a disease phenotype - dilated cardiomyopathy. We extracted from the DisGeNET database (25) the list of genes associated with cardiomyopathies and overlapped it with the list of genes with AS events found only at 200M. We found two low-expressed (TPM < 1) genes with AS that might be associated with cardiomyopathies: FKBP1B (DisGeNet score 0.2) and CDH23 (DisGeNet score 0.1). We further confirmed the AS events in these genes by visualizing the read coverage via the sashimi tool (24). Figure 4 demonstrates that the AS events exist in both genes and are consistent between all heart (DCM) samples.

**Figure 4.**
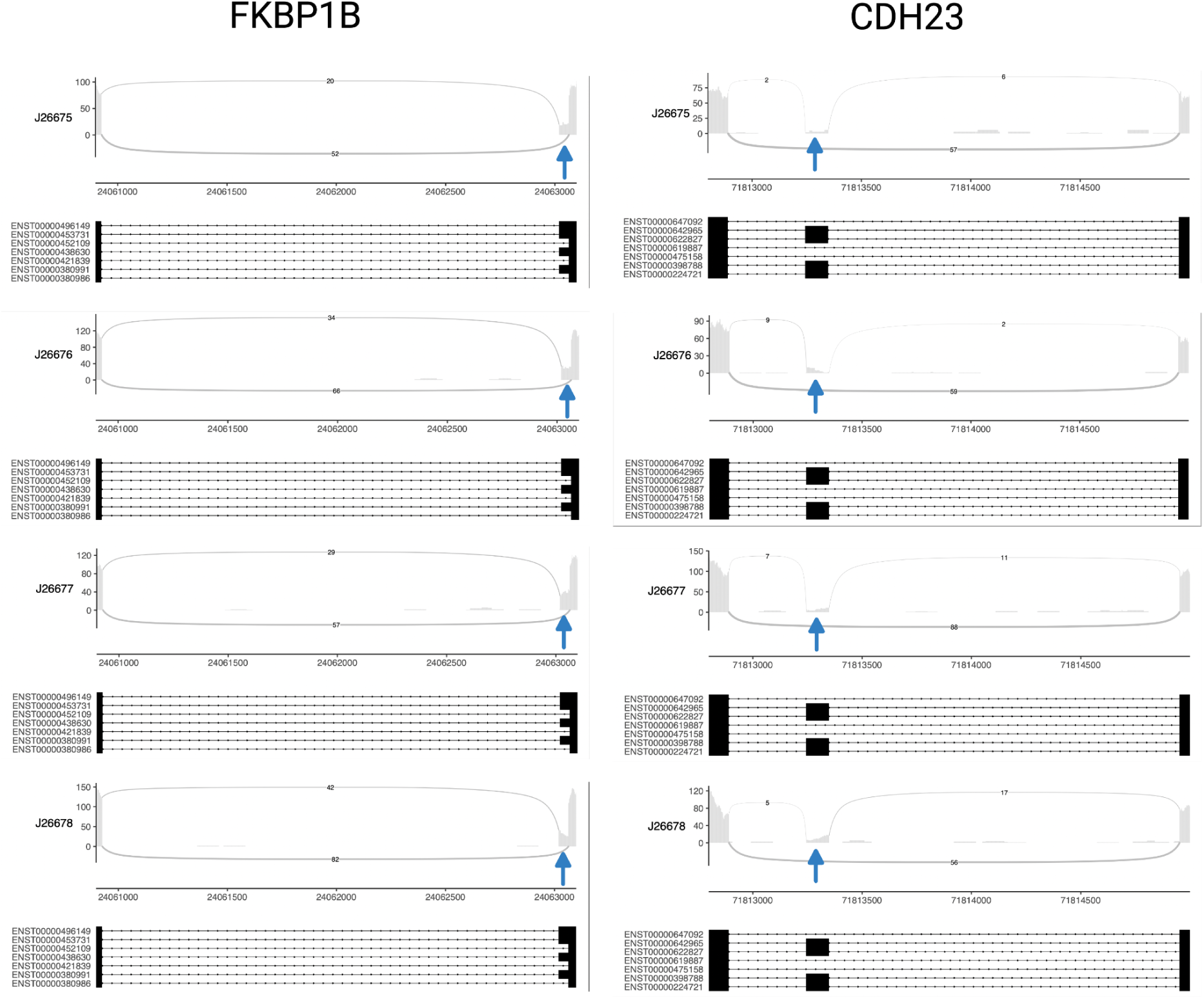
Sashimi plots for low expressed genes FKBP1B and CDH23 with AS detected only at 200M reads. The blues arrows indicate alternative splice site (FKBP1B) and exon skipping (CDH23) events.

Finally, we performed AS-aware functional enrichment analysis using NEASE (23) with KEGG pathways on the exon skipping events detected at 50M, 100M, 150M, and 200M. We extracted the pathways that could be detected only at 200M reads (Table S7). Among KEGG pathways for the adipose sample before the treatment, we found “Fatty acid degradation” (p-adjusted 0.043). For the endotoxin-treated adipose sample, we found the response to viral infections (p-adjusted 0.01). Finally, for the hypothalamus sample, the KEGG pathways list contains terms related to the neural system such as ‘Gap junction’ (p-adjusted 0.01), Adherens junction (p-adjusted 0.02), and Glutamatergic synapse (p-adjusted 0.02).

Altogether, the analysis of genes with AS that can be detected only at 200M reads reveals that the biologically relevant signals might be weak but still present and worth investigating for a more comprehensive view.

### Comparison of GTEx and TCGA cohorts with the deep-sequenced cohorts

The GTEx (15) and TCGA cohorts comprise a large amount of RNA-Seq data from healthy individuals and patients with cancer, respectively. The usual sequencing depth for those cohorts varies between 50-150M reads. As we showed on the SARS-CoV-2 dataset, such sequencing depth might not be enough for a comprehensive AS analysis. We compared the number of genes/junctions with AS we can recover from the GTEx and TCGA RNA-Seq data with the amount that could be recovered with the higher sequencing depth estimated from the deep-sequenced cohorts. For the estimation, we used the most simple assumption that the number of events would increase linearly or quadratically.

For the GTEx cohort analysis, we used 102 samples from the hypothalamus with a sequencing depth of 10-236M reads. We detected AS in these samples and used the number of detections to estimate the number of genes/junctions involved in AS for higher sequencing depth (Figure 5) using a polynomial fit of degree 1 (blue dashed line) and 2 (orange dashed line). We also approximated these values based on the number of genes/junctions involved in AS detected in samples after downsampling the hypothalamus sample with 236M reads disregarding the gene expression level (green dashed line). The resulting number of detected genes and junctions has also been used in a polynomial fit of a degree 2.

**Figure 5:**
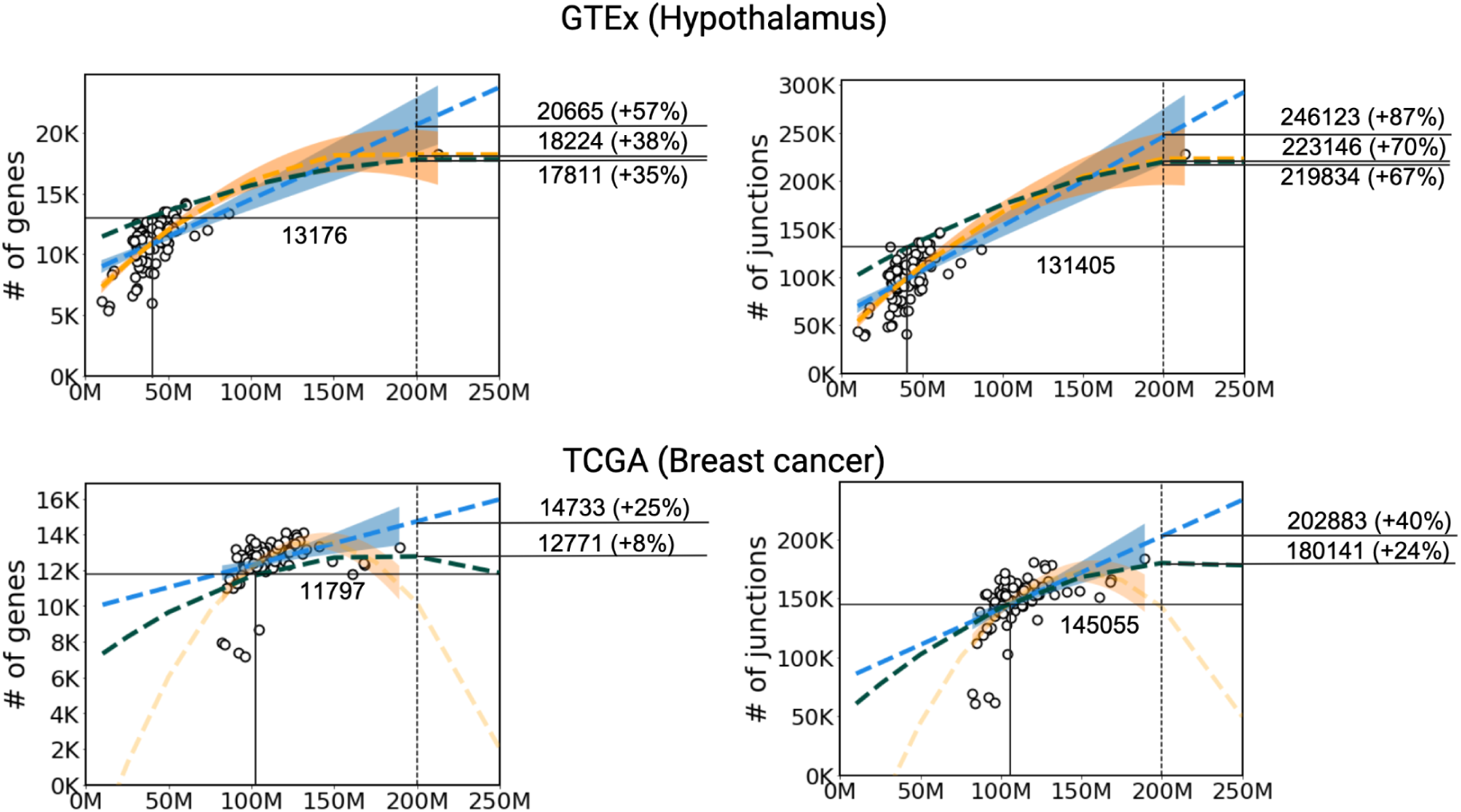
The real (dots) and estimated (dashed lines) number of genes with AS (left) and junctions involved in AS (right) in GTEx hypothalamus and TCGA breast cancer data sets. The number indicates the number of detections at a baseline (black horizontal line), estimated from the downsampling analysis (green dashed line), and estimated from the real datasets using the polynomial fit with the degrees of freedom equal to 1 (blue dashed line) and 2 (orange dashed line). For the percentage, the number of detections in a baseline corresponds to 100%. The shaded area indicates a confidence interval.

**Figure 6:**
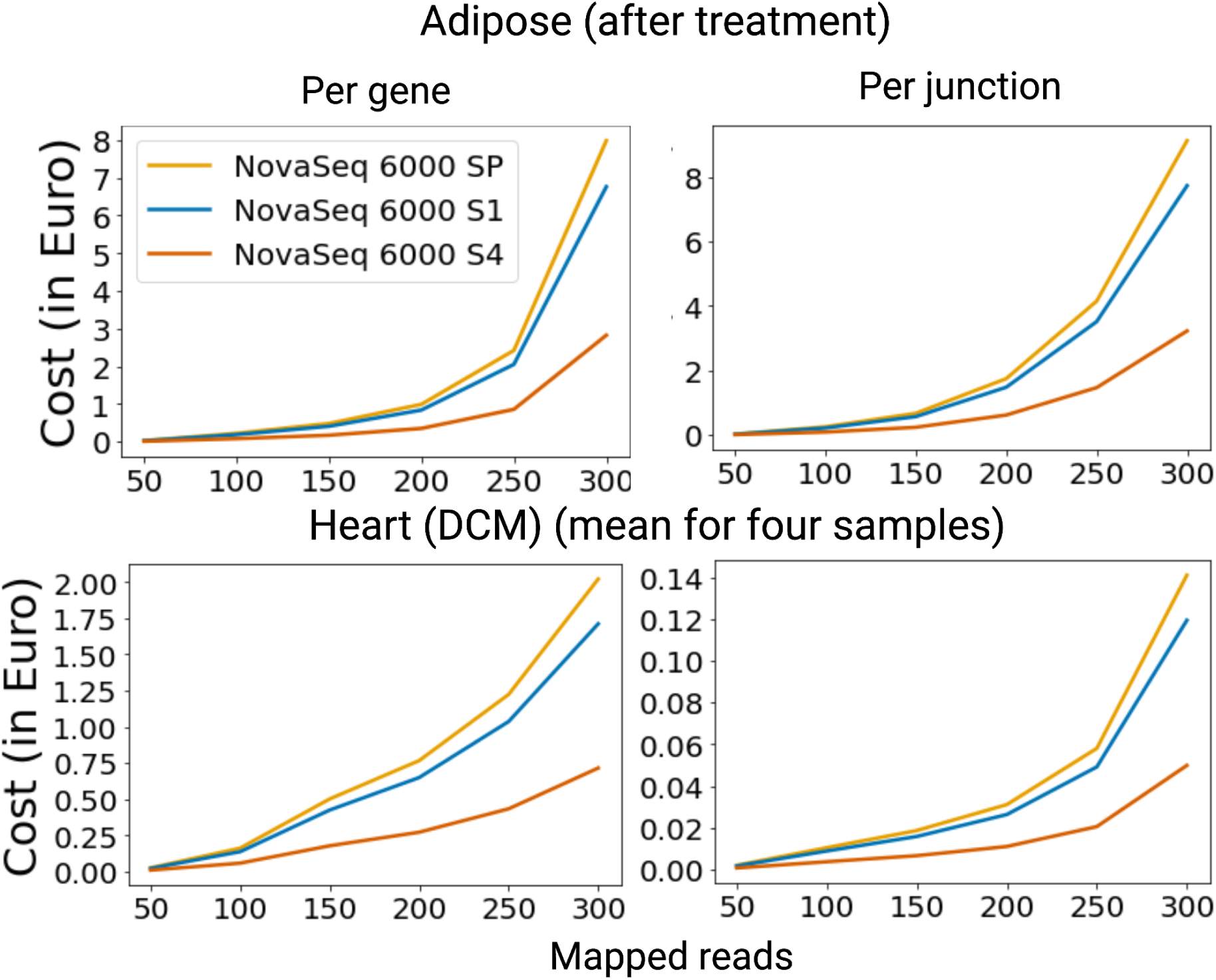
The cost for each additional gene with AS and junction detected while increasing the number of mapped reads

Figure 5 (top) demonstrates the possible loss in AS detection in the GTeX dataset. As the baseline (black lines), we used the number of genes/junctions that would be detected in a sample with a sequencing depth equal to the mean sequencing depth of the Hypothalamus GTEx dataset of 41 M. Analysis of a sample with the mean sequencing depth would result in the possible loss of from up to 7489 genes with AS/114717 alternative junctions.

For the TCGA breast cancer analysis, we randomly selected 100 samples from the breast cancer cohort and detected genes with AS and alternative junctions. We estimated the number of genes with AS/alternative junctions for higher sequencing depth using a polynomial fit of degree 1 (blue dashed line) and 2 (orange dashed line). Unfortunately, the polynomial fit of degree 2 failed to approximate the number of genes/junctions involved in AS as there were no samples of more than 200M reads. Thus, for the TCGA dataset, we performed the downsampling analysis similar to the analysis of deep-sequenced data sets using the sample with the maximum number of reads found in this part of the cohort equal to 189M. We used the results of the downsampling to approximate the increase in the number of genes/ junctions involved in AS with the increase of sequencing depth by the polynomial that fits a degree of 2 (green dashed line).

Figure 5 (bottom) demonstrates the possible loss in AS detection in the TCGA breast cancer cohort. As the baseline (horizontal black line), we also used the number of genes/junctions that would be detected in a sample with a sequencing depth equal to the mean sequencing depth of 109M. Analysis of a sample with the mean sequencing depth would result in the possible loss of up to 2935 genes with AS/57827 alternative junctions.

Summarizing, both of these RNA-seq datasets are undersequenced for comprehensive AS analysis, especially the samples in the GTEx cohort.

### The cost of AS detection in the deep-sequenced cohorts

Experiments with more than 200M reads might be economically impractical. We used the prices published on the CCGA Kiel website and estimated/calculated that the cost for each additional detection (a gene or a junction with AS) grows notably after 200-250M reads (Figure 5). Together with the results from deep-sequenced cohorts, this result supports the threshold of 200M reads as an optimal trade-off between costs and benefits for AS analysis.

## DISCUSSION

The appropriate choice of sequencing depth is crucial for successful RNA-Seq analysis, especially when studying alternative splicing. Our results demonstrate that the threshold depends on the purpose of the study: to detect all AS events, the threshold should be set to 150M-200M reads for lowly expressed genes and 100-150M for highly expressed genes. Thus, we observed the increase of new AS detections to be less than 1 % after these thresholds.

Several aspects need to be considered when interpreting this threshold. The sequencing depth of a sample is typically higher than the number of mapped reads, however, in high-quality experiments, those numbers should be close to each other. We defined the threshold based on short-read paired-end mRNA RNA-Seq data from human, and we should note that experimental design might impact the results. First, the choice of sequencing technology and library preparation affects this threshold. For example, single-end short sequencing data which is less robust for gene and isoform expression analysis [7] likely has different thresholds. Read length and the level of RNA degradation might also impact the threshold for alternative splicing detection: shorter length decreases the fraction of uniquely mapped reads. Concerning RNA degradation, in our study, we did not address its effect on alternative splicing detection, as the analyzed datasets present high-quality data (the RIN (RNA integrity number) value 8-10), and the datasets with the RIN value less than 7 is generally not recommended to use for the transcriptome analysis.

Similarly, we expect this threshold to differ for long-read sequencing techniques which can detect the full-length and novel transcripts. However, despite the rapid development of long-read sequencing technology, like the increase in base calling and transcript identification accuracy (26), this technology is currently more expensive and requires the extraction of a larger amount of undamaged nucleotide material which might be challenging. Thus short-read sequencing technologies are currently more widely used for quantitative alternative splicing analysis and present the best option in terms of cost-effectiveness and requirements to the material preparation.

Another limitation of this study is the selection of the organism: all found deep-sequenced RNA-Seq data sets have been derived from the human samples. For other organisms, the threshold might differ because of the difference in splicing patterns, e.g., in humans, exon skipping was shown to be the prevalent alternative splicing event [9] but in plants, intron retention prevails [10]. However, the threshold for the sequencing depth defined in our study is robust regardless of disease or tissue type.

Finally, our results may be affected by the computational workflow selected. While the mapping tool should not impact the results [11], choosing an alternative splicing detection tool might impact the threshold as the number of detected alternative splicing events differs notably between tools [12]. However, a previous study on human adipose tissue with a different alternative splicing detection tool - MATS - demonstrated similar results [3].

Before choosing the number of reads for a particular experiment, one more important question is how relevant alternative splicing events detected only at deeper sequencing could be to the phenotype/disease of interest. Deveson et al (27) demonstrated that with increasing sequencing depth noncoding introns and isoforms also kept increasing. However, the event level resolution we use in the study cannot consider if the resulting isoform will be coding or not. On the gene level, we do not see any prevalence of alternative splicing events in the non-protein-coding genes (data not shown). Our study demonstrated that the higher sequencing depth results in more alternative splicing events in genes with low and high (but not very high) expression levels. Each of these events brings only a subtle change. Still, if they accumulate in one metabolic pathway or protein complex, the additive effect might lead to the phenotype/disease onset. E.g., we demonstrated for the hypothalamus and adipose datasets, that the changes detected at higher sequencing depth accumulate in phenotype-specific pathways. Our results suggest that the threshold of 150-200M reads for lowly and 100-150M highly expressed genes present a trade-off between the number of detections to reveal the role of alternative splicing in the phenotype/disease of interest and cost-effectiveness. Thus, the typical threshold for sequencing depth chosen in contemporary human RNA-Seq studies e.g., TCGA and GTEx is too low for a complete assessment of the alternative splicing landscape, and future projects may consider sequencing deeper to better support alternative splicing analysis.

## DATA AVAILABILITY

The preprocessed data underlying this article are available in Zenodo at https://doi.org/10.5281/zenodo.11655945. The GTEx data underlying this article were obtained from dbGaP accession number phs000424.GTEx.v8.p2 on 21/02/2020. The TCGA data used here are, in part, based upon data generated by the TCGA Research Network: https://www.cancer.gov/tcga. The SARS-CoV-2 and human adipose RNA-Seq data are available in Gene Expression Omnibus (GEO) (14) at https://www.ncbi.nlm.nih.gov/Traces/study/?acc=PRJNA787951 and https://www.ncbi.xyz/Traces/study/?acc=PRJNA198737 and can be accessed with GSE190680, GSE46323. The Heart (DCM) data will be shared on reasonable request to the corresponding author. All Python scripts used to generate the figures are available at https://github.com/OlgaVT/RNA-Seq Depth for AS.

## AUTHOR CONTRIBUTIONS

Olga Tsoy: Conceptualization, Methodology, Analysis, Writing - original draft. Sabine Ameling: RNA-Seq data from the heart tissues of patients with DCM, Writing - review and editing. Sören Franzenburg: RNA-Seq data from the heart tissues of patients with DCM, Writing - review and editing. Markus Hoffmann: the preprocessing of the SARS-Cov2 and TCGA RNA-Seq data, Writing - review and editing. Lina Liv-Willuth: the preprocessing of the SARS-Cov2 RNA-Seq data, Writing - review and editing. Hye Kyung Lee: the SARS-Cov2 RNA-Seq data, Writing - review and editing. Ludwig Knabl: the SARS-Cov2 RNA-Seq data, Writing - review and editing. Priscilla A. Furth: the SARS-Cov2 RNA-Seq data, Writing - review and editing. Uwe Völker: Conceptualization, RNA-Seq data from the heart tissues of patients with DCM, Writing - review and editing. Lothar Hennighausen: the SARS-Cov2 RNA-Seq data, Writing - review and editing. Jan Baumbach: Conceptualization, Methodology, Writing - review and editing. Tim Kacprowski: Conceptualization, Methodology, Writing - review and editing. Markus List: Conceptualization, Methodology, Access to the RNA-Seq data from the TCGA cohort, Writing - review and editing.

