## Supplementary Material for "RNA sequencing depth guidelines for the study of alternative splicing"

### KAT5

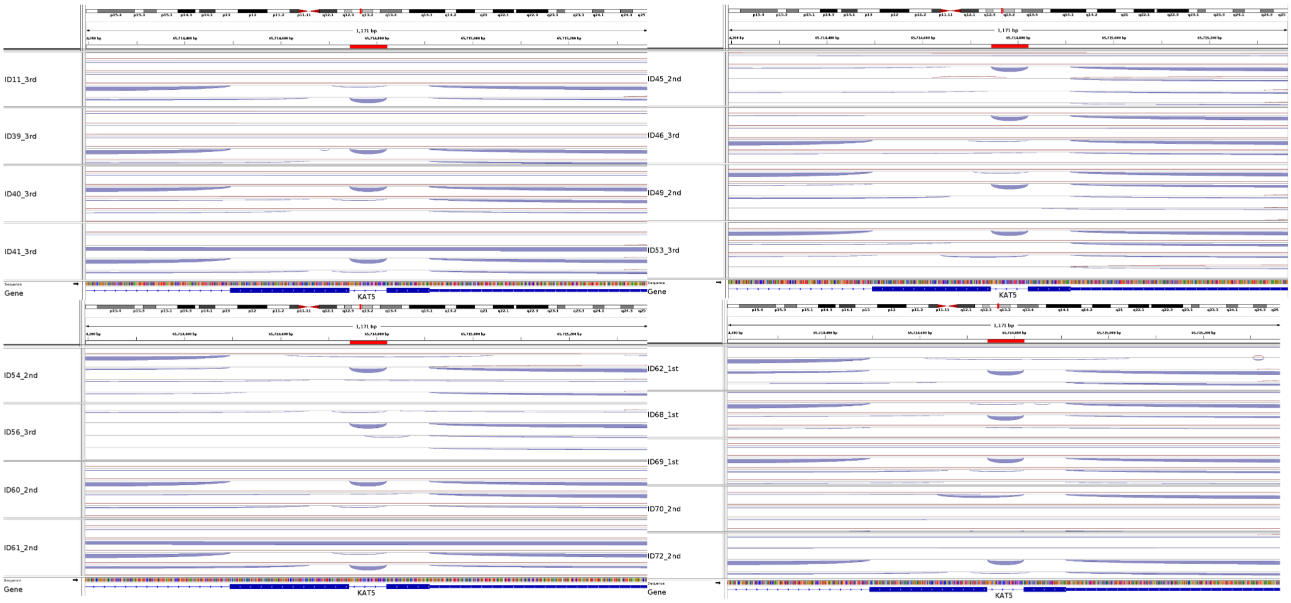

ID11_3rd
ID39_3rd
ID40_3rd
ID41_3rd
Gene
KAT5
ID45_2nd
ID46_3rd
ID49_2nd
ID53_3rd
Gene
KAT5
ID54_2nd
ID56_3rd
ID60_2nd
ID61_2nd
Gene
KAT5
ID62_1st
ID68_1st
ID69_1st
ID70_2nd
ID72_2nd
Gene
KAT5


### KIF21A

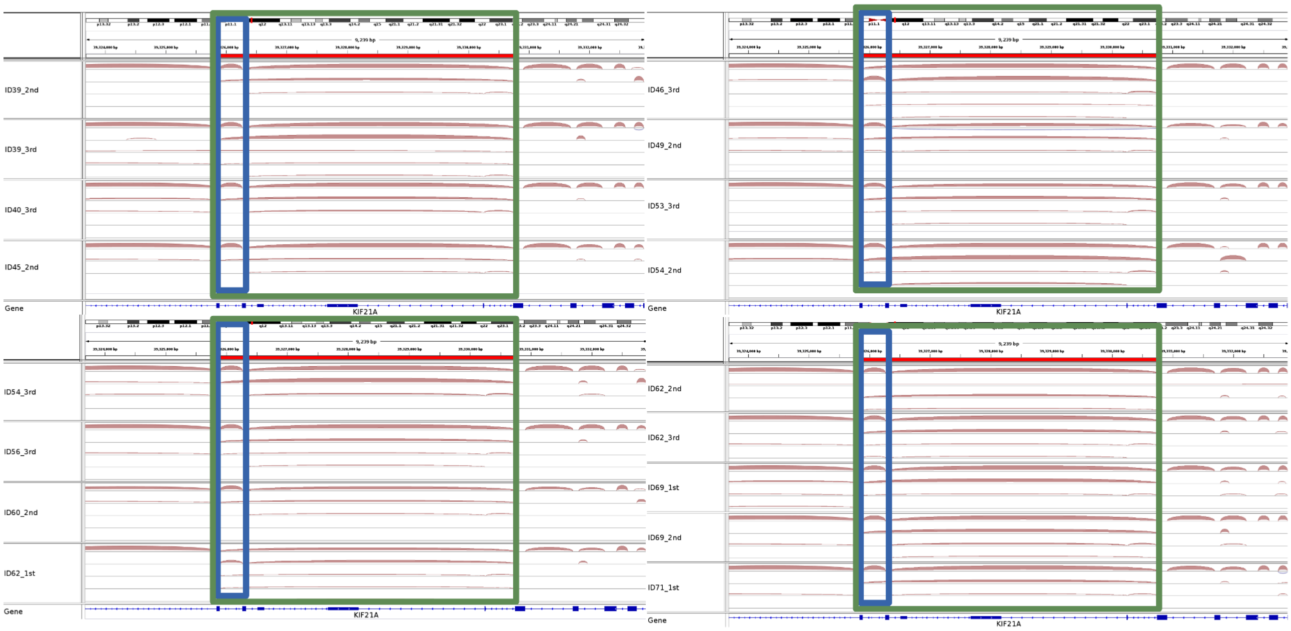

ID39_2nd
ID39_3rd
ID40_3rd
ID45_2nd
Gene
KIF21A
ID46_3rd
ID49_2nd
ID53_3rd
ID54_2nd
Gene
KIF21A
ID54_3rd
ID56_3rd
ID60_2nd
ID62_1st
Gene
KIF21A
ID62_2nd
ID62_3rd
ID69_1st
ID69_2nd
ID71_1st
Gene
KIF21A

# NR6A1

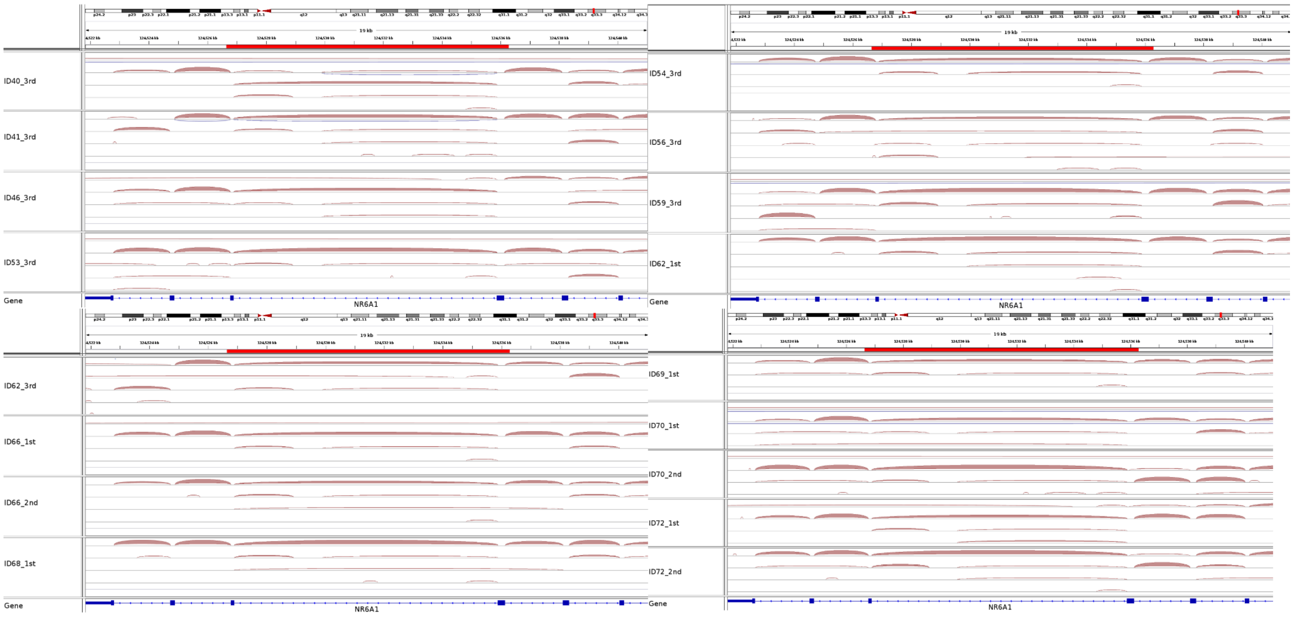


### PRKACB

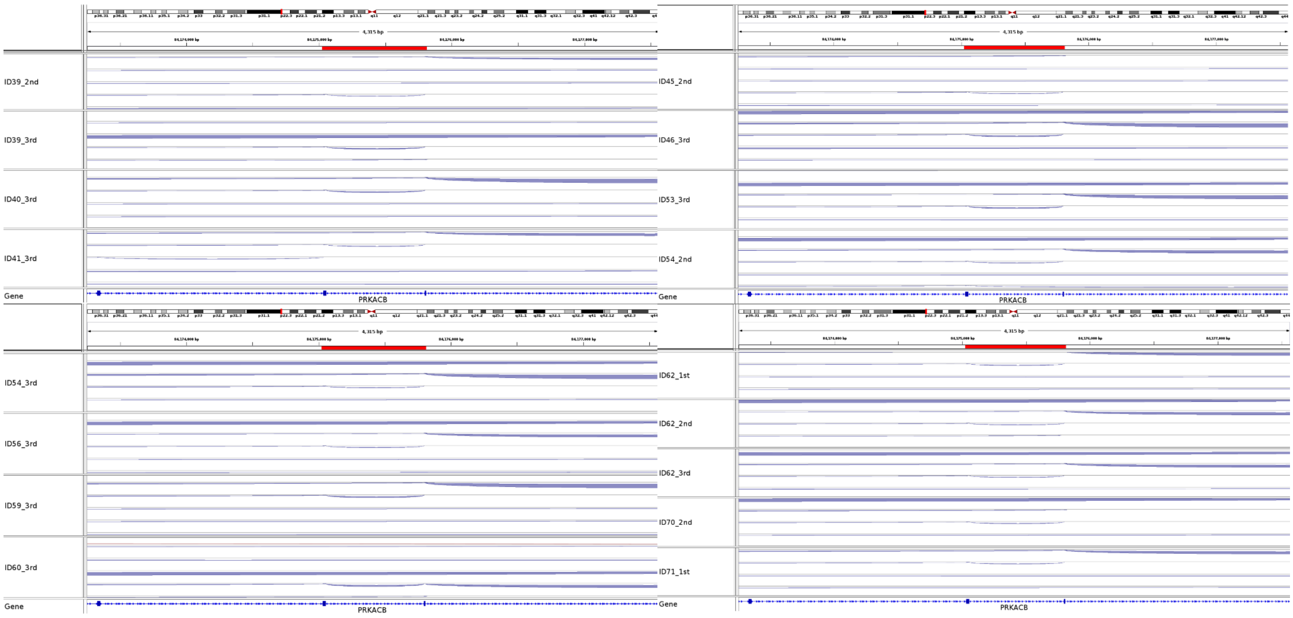

## TP53BP2

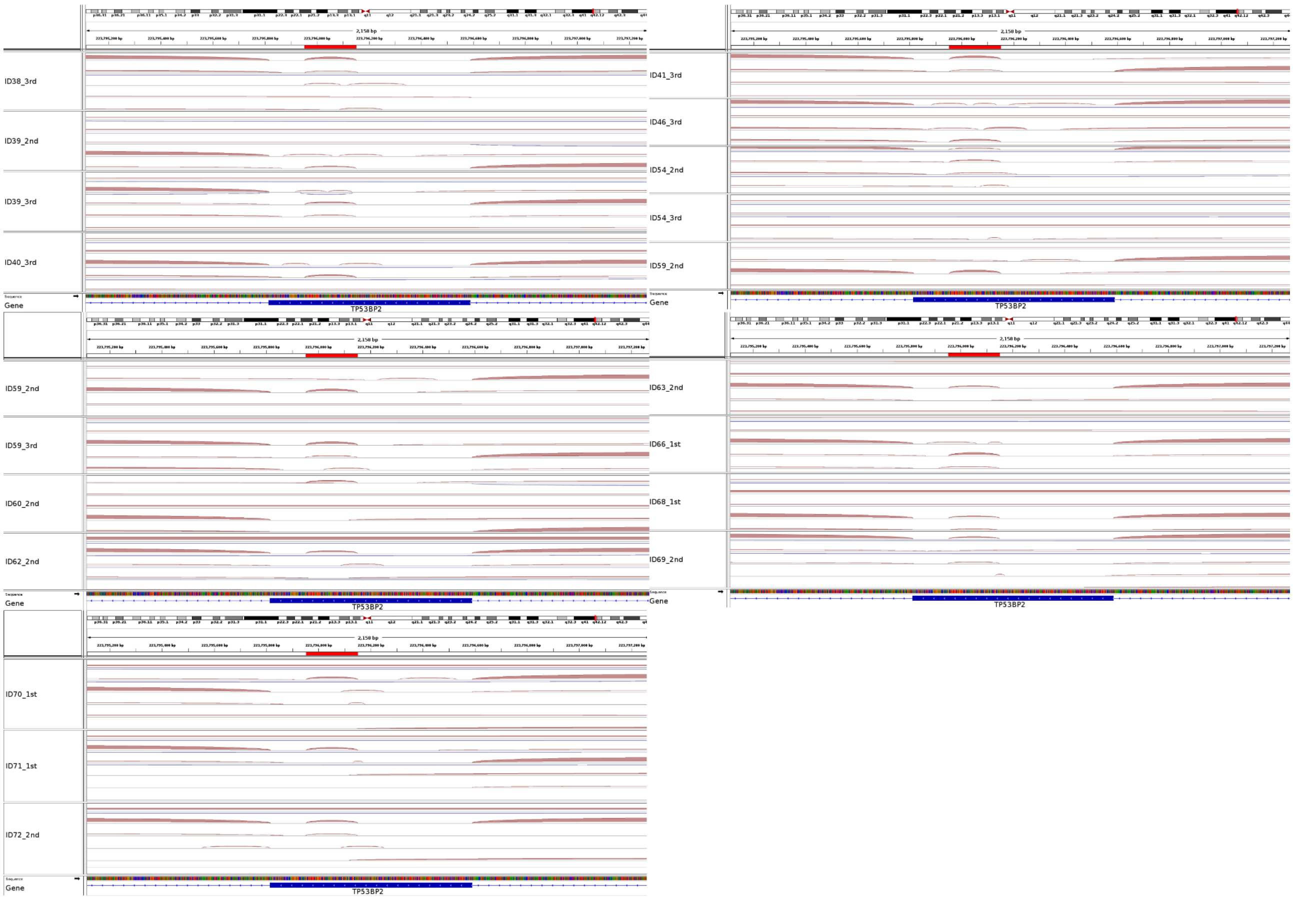


#### ZEB2

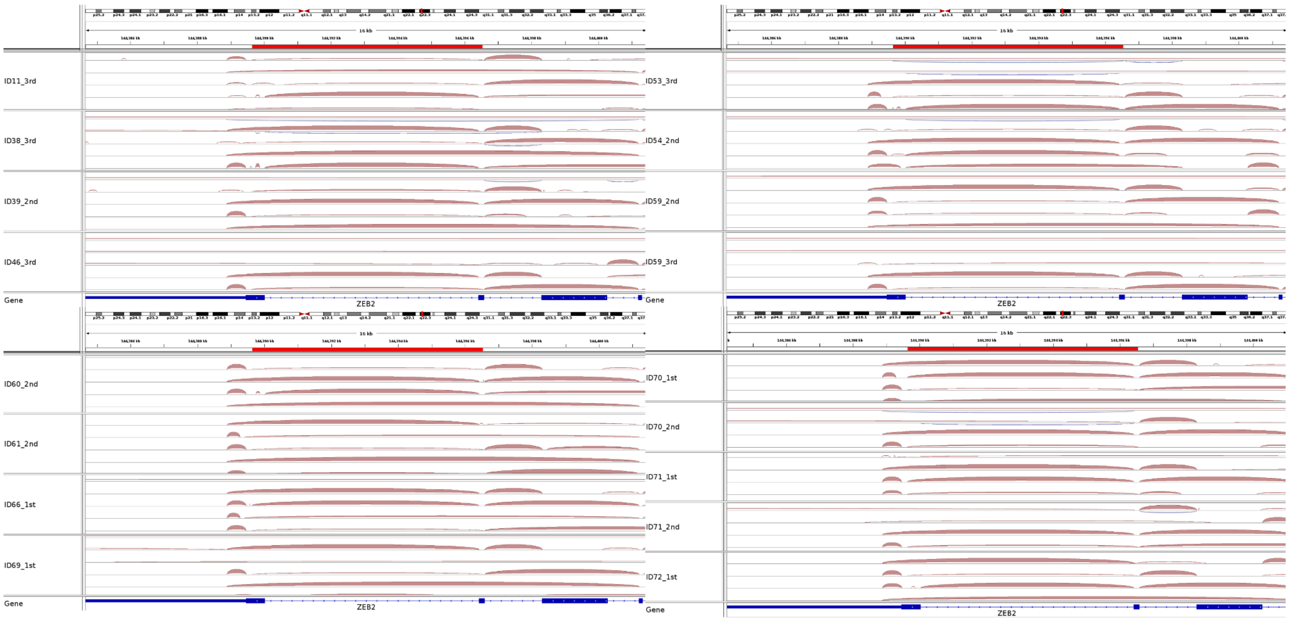


**Supplementary Figure S1.** Read coverage of junctions detected to be involved in AS only in samples with a higher sequencing depth.

The red line marks the coordinates of the alternative junction and the plots below represent the coverage that supports these junctions from the alignment bam files per each sample.

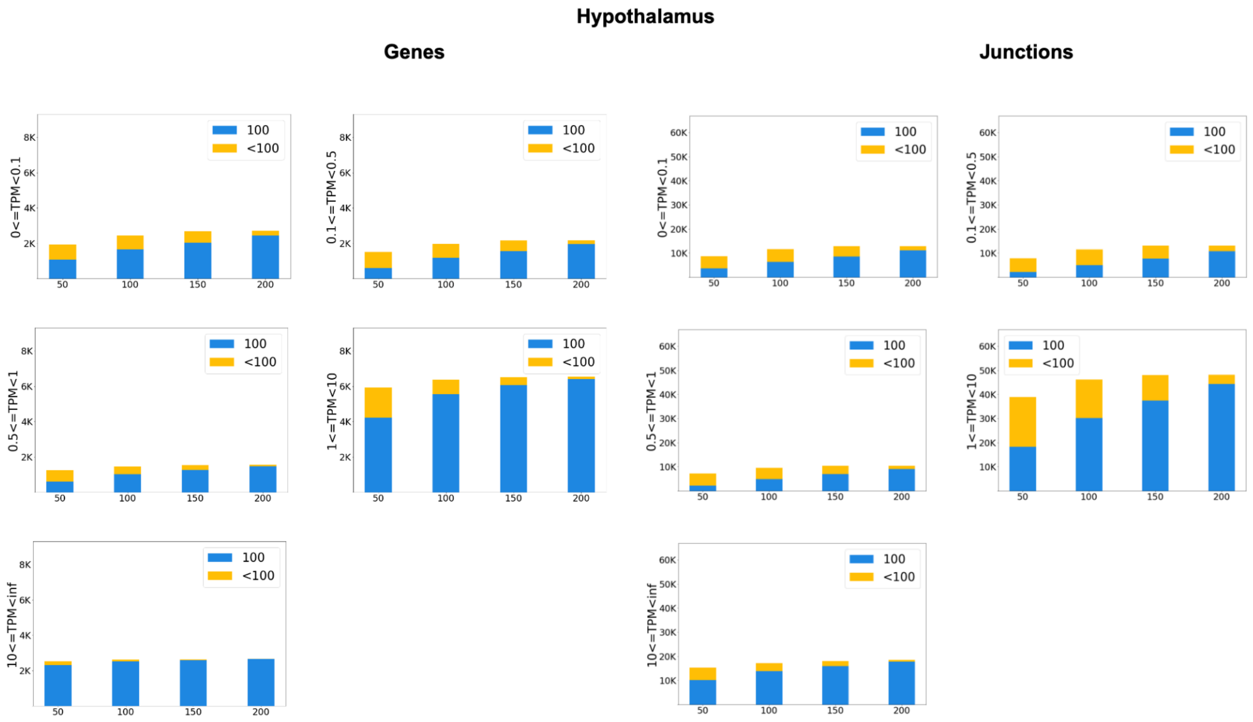


**Supplementary Figure S2.** The number of genes and junctions involved in AS detected in 100 (blue), and less than 100 (yellow) out of 100 subsamples for Hypothalamus. Each stacked bar depicts one sample.

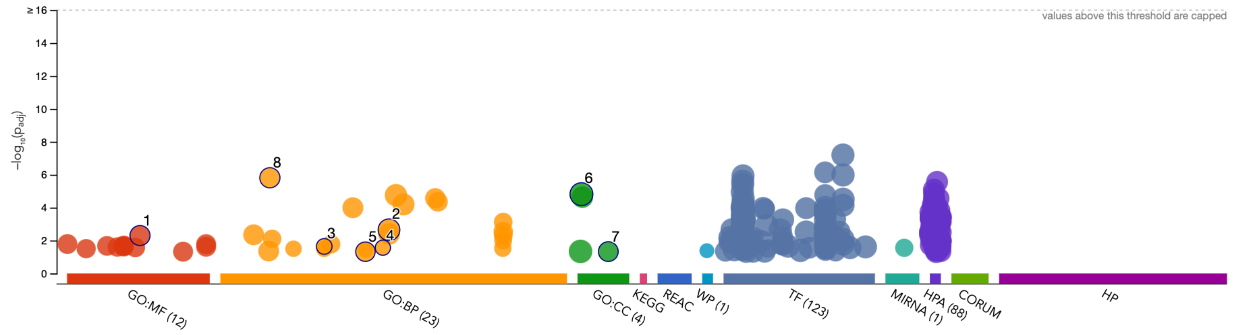


| ID | Source | Term ID | Term Name | $p_{adj}$ (query_1) |
|---|---|---|---|---|
| 8 | GO:BP | GO:0010033 | response to organic substance | $1.532\times10^{-6}$ |
| 6 | GO:CC | GO:0005737 | cytoplasm | $1.517\times10^{-5}$ |
| 2 | GO:BP | GO:0048518 | positive regulation of biological process | $2.210\times10^{-3}$ |
| 1 | GO:MF | GO:0043168 | anion binding | $4.950\times10^{-3}$ |
| 3 | GO:BP | GO:0032869 | cellular response to insulin stimulus | $2.269\times10^{-2}$ |
| 4 | GO:BP | GO:0046718 | symbiont entry into host cell | $2.714\times10^{-2}$ |
| 7 | GO:CC | GO:0070062 | extracellular exosome | $4.674\times10^{-2}$ |
| 5 | GO:BP | GO:0044419 | biological process involved in interspecies inte... | $4.748\times10^{-2}$ |

**Supplementary Figure S3.** The results of the gProfiler for the genes with high expression (TPM>1) with AS detected only at 200M reads from the adipose sample after endotoxin treatment.

**Supplementary Table S1.** Alternative splicing tools that simulated RNA-Seq data at various sequencing depths for the evaluation of the tool performance

| Tool name | Maximum sequencing depth of the simulated RNA-Seq data | Reference |
|---|---|---|
| ASpedia-FI | 52M | (1) |
| AS-Quant | 50M | (2) |
| ASTool | 60M | (3) |
| iREAD | 30M | (4) |
| splAdder | 20 M for 1000 genes | (5) |
| McSplicer | 75M | (6) |
| FreePSI | 50M | (7) |
| SUPPA2 | 120M | (8) |
| NURD | 10M | (9) |
| Cufflinks | 100M | (10) |
| CASH | 100M | (11) |
| EventPointer3.0 | 125M | (12) |
| ASimulatoR | 200M | (13) |
| DICAST | 200M | (14) |

**Supplementary Table S2.** Number of genes and junctions in each expression category

| | SARS-Cov2 | Hypothalamus | Adipose (pre-treatment) | Adipose (post-treatment) | Heart (DCM) |
|---|---|---|---|---|---|
| | **Genes** | | | | |
| TPM < 0.1 | 6149 ± 647 | 10303 | 11863 | 12010 | 13436 ± 412 |
| 0.5 > TPM ≥0.1 | 4213 ± 158 | 6179 | 5394 | 5249 | 4889 ± 98 |
| 1 > TPM ≥ 0.5 | 2006 ± 63 | 2825 | 2380 | 2204 | 2187 ± 53 |
| 10 > TPM ≥ 1 | 7254 ± 281 | 8771 | 5568 | 4586 | 4967 ± 229 |
| TPM ≥ 10 | 5112 ± 638 | 3203 | 1959 | 1580 | 1307 + 88 |
| | **Junctions** | | | | |
| TPM < 0.1 | 24302 ± 3980 | 55356 | 54227 | 63026 | 107673 ± 1855 |

| | | | | | |
|---|---|---|---|---|---|
| 0.5 > TPM ≥0.1 | 32095 ± 3980 | 70978 | 58738 | 68798 | 95009 ± 1158 |
| 1 > TPM ≥ 0.5 | 21823 ± 3106 | 45310 | 28151 | 38621 | 52799 ± 1620 |
| 10 > TPM ≥ 1 | 144588 ± 18131 | 178510 | 63872 | 101345 | 120282 ± 7253 |
| TPM ≥ 10 | 154833 ± 23376 | 66807 | 26405 | 40517 | 29891 ± 2066 |

**Supplementary Table S3**. The value of slopes from the significant results of the Mann-Kendall Trend test (p-value < 0.05).

| | **Heart (DCM) Sample 1** | **Heart (DCM) Sample 2** | **Heart (DCM) Sample 3** | **Heart (DCM) Sample 4** | **Adipose (pre-treatment)** | **Adipose (post-treatment)** |
|---|---|---|---|---|---|---|
| | Genes (In at least one subsample) | | | | | |
| TPM < 0.1 | 304.67 | 310.67 | 291.33 | 325.0 | 165.33 | No trend |
| 0.5 > TPM ≥ 0.1 | 41.67 | 66.33 | 57.33 | 67.33 | No trend | No trend |
| 1 > TPM ≥ 0.5 | 11.0 | 15.33 | 27.33 | 27.33 | 29 | 16.67 |
| 10 > TPM ≥ 1 | 44.0 | 39.4 | 38.0 | 42.33 | 49 | No trend |
| TPM ≥ 10 | 25.0 | 28.0 | 27.0 | 32.0 | 22 | 4.33 |
| | Genes (In 100 subsamples) | | | | | |
| TPM < 0.1 | 394.0 | 379.67 | 377.0 | 376.0 | 294.33 | 240.33 |
| 0.5 > TPM ≥ 0.1 | 236.67 | 245.33 | 237.67 | 232.0 | 294.67 | 224.67 |
| 1 > TPM ≥ 0.5 | 82.0 | 90.33 | 89.33 | 87.67 | 95 | 86.67 |
| 10 > TPM ≥ 1 | 129.33 | 132.0 | 136.0 | 139.0 | 195.33 | 123.67 |
| TPM ≥ 10 | 40.67 | 43.33 | 45.0 | 51.0 | 47.33 | 39 |
| | Junctions (In at least one subsample) | | | | | |
| TPM < 0.1 | 2188.33 | 2141.33 | 2029.33 | 2099.67 | 979 | No trend |
| 0.5 > TPM ≥ 0.1 | No trend | No trend | 752.67 | 801.67 | No trend | No trend |

| | | | | | | |
|---|---|---|---|---|---|---|
| 1 > TPM ≥ 0.5 | No trend | No trend | No trend | No trend | 823.33 | 378 |
| 10 > TPM ≥ 1 | 162.0 | 150.67 | 109.0 | 216.0 | No trend | No trend |
| TPM ≥ 10 | 158.33 | 173.33 | 177.0 | 191.33 | 251 | No trend |
| | Junctions (In 100 subsamples) | | | | | |
| TPM < 0.1 | 1933.0 | 1925.4 | 1834.0 | 1758.0 | 1384.33 | 937 |
| 0.5 > TPM ≥ 0.1 | 1865.0 | 1851.67 | 1817.0 | 1770.33 | 1851.33 | 1238 |
| 1 > TPM ≥ 0.5 | 625.0 | 731.0 | 731.33 | 682.33 | 884.33 | 1365 |
| 10 > TPM ≥ 1 | 932.0 | 1059.0 | 1059.0 | 1052.0 | 1989.67 | 902.67 |
| TPM ≥ 10 | 204.0 | 228.0 | 223.0 | 230.33 | 470.67 | 263 |

**Supplementary Table S4.** The increase of the mean number and the percentage of genes and junctions involved in alternative splicing with increasing sequencing depth in the Adipose (pre-treatment) dataset. Column '50 M' indicates the mean number and the percentage of genes/junctions detected in the samples with a sequencing depth of less than 50 M reads. The column '300M' indicates the mean number and the percentage of genes/junctions detected in the samples with the highest sequencing depth of 300M reads. Grey shading indicates an increase of less than 1% of new detections.

| TPM | 50M | 50-100 | 100-150 | 150-200 | 200-250 | 250-300 | 300M | Total # of genes/ junctions |
|---|---|---|---|---|---|---|---|---|
| | Genes (In at least one subsample) | | | | | | | |
| < 0.1 | 2005<br>(16.9%) | +635<br>(+5.4%) | +324<br>(+2.7%) | +136<br>(+1.1%) | +36<br>(+0.3%) | -24<br>(-0.2%) | 3112<br>(26.2%) | 11863 |
| [0.1, 0.5) | 1970<br>(36.5%%) | +460<br>(+8.5%%) | +176<br>(+3.2%%) | +65<br>(+1.2%%) | -7<br>(-0.1%%) | -14<br>(-0.3%) | 2650<br>(49.1%%) | 5394 |
| [0.5, 1) | 1286<br>(54.0%%) | +185<br>(+7.8%) | 62<br>(+2.6%) | 15<br>(+0.6%) | 8<br>(+0.3%) | -2<br>(-0.08%) | 1554<br>(65.3%%) | 2380 |
| [1, 10) | 3765<br>(67.6%) | +202<br>(+3.6%) | +94<br>(+1.7%) | +40<br>(+0.7%) | +13<br>(+0.2%) | +8<br>(+0.1%) | 4122<br>(74.0%) | 5568 |

| | | | | | | | | |
|---|---|---|---|---|---|---|---|---|
| ≥ 10 | 1381 (70.5%%) | 94 (+4.7%) | 39 (+2.0%) | 18 (+0.9%) | 9 (+0.5%) | 5 (+0.3%) | 1546 (78.9%) | 1959 |
| | **Genes (In 100 subsamples)** | | | | | | | |
| < 0.1 | 858 (7.2%) | 495 (+4.2%) | 346 (+2.9%) | 273 (+2.3%) | 264 (+2.2%) | 248 (+2.1%) | 2484 (20.9%) | 11863 |
| [0.1, 0.5) | 703 (13.0%) | 569 (+10.5%) | 384 (+7.1%) | 279 (+5.2%) | 221 (+4.1%) | 167 (+3.1%) | 2323 (43.0%) | 5394 |
| [0.5, 1) | 620 (26.1%) | 383 (+16.1%) | 171 (+7.2%) | 122 (+5.1%) | 75 (+3.2%) | 66 (+2.8%) | 1437 (60.4%) | 2380 |
| [1, 10) | 2591 (46.5%) | 709 (+12.7%) | 310 (+5.6%) | 159 (+2.9%) | 117 (+2.1%) | 89 (+1.6%) | 3975 (71.4%) | 5568 |
| ≥ 10 | 1225 (62.5%) | 137 (+7.0%) | 78 (+4.0%) | 40 (+2.0%) | 24 (+1.2%) | 21 (+1.1%) | 1525 (77.8%) | 1959 |
| | Junctions (In at least one subsample) | | | | | | | |
| < 0.1 | 7472 (13.8%) | +3345 (+6.2%) | +1832 (+3.4%) | +928 (+1.5%) | +276 (+0.5%) | –8 (–0.01%) | 13746 (25.3%) | 54227 |
| [0.1, 0.5) | 8810 (15.0%) | +4448 (+7.6%) | +2028 (+3.5%) | +634 (+1.1%) | +90 (+0.2%) | –130 (–0.2%) | 15880 (27.0%) | 58738 |
| [0.5, 1) | 6446 (22.9%) | +2215 (+7.9%) | +883 (+3.1%) | +228 (+0.8%) | –89 (–0.3%) | –85 (+0.3%) | 9598 (34.1%) | 28151 |
| [1, 10) | 19043 (29.8%) | +3399 (+5.3%) | +956 (+1.5%) | +80 (+0.1%) | –135 (–0.2%) | –22 (–0.03%) | 23321 (36.5%) | 63872 |
| ≥ 10 | 5539 (21.0%) | +782 (+3.0%) | +370 (+1.4%) | +250 (+0.9%) | +133 (+0.5%) | +157 (+0.6%) | 7231 (27.4%) | 26405 |
| | Junctions (In 100 subsamples) | | | | | | | |
| < 0.1 | 2502 (4.6%) | +1831 (+3.4%) | +1520 (+2.8%) | +1317 (+2.4%) | +1316 (+2.4%) | +1376 (+2.5%) | 9862 (18.2%) | 54227 |
| [0.1, 0.5) | 2117 (2.6%) | +2507 (+4.2%) | +2170 (+3.7%) | +1724 (+2.9%) | +1660 (+2.8%) | +1700 (+2.9%) | 11878 (20.2%) | 58738 |
| [0.5, 1) | 2018 (7.2%) | +1789 (+6.4%) | +1280 (+4.5%) | +957 (+3.4%) | +841 (+3.0%) | +790 (+2.8%) | 7675 (27.3%) | 28151 |
| [1, 10) | 9075 (14.2%) | +4216 (+6.6%) | +2661 (+4.2%) | +1812 (+2.8%) | +1496 (+2.3%) | +1369 (+2.1%) | 20629 (32.3%) | 63872 |
| ≥ 10 | 4089 (15.5%) | +953 (+3.6%) | +623 (+2.4%) | +430 (+1.6%) | +359 (+1.4%) | +352 (+1.3%) | 6806 (25.8%) | 26405 |

**Supplementary Table S5.** The increase of the mean number and the percentage of genes and junctions involved in alternative splicing with increasing sequencing depth in the Adipose (post-treatment) dataset. Column '50 M' indicates the mean number and the percentage of genes/junctions detected in the samples with a sequencing depth of less than 50 M reads. The column '300M' indicates the mean number and the percentage of genes/junctions detected in the samples with the highest sequencing depth of 300M reads. Grey shading indicates an increase of less than 1% of new detections.

| TPM | 50M | 50-100 | 100-150 | 150-200 | 200-250 | 250-300 | 300M | |
|---|---|---|---|---|---|---|---|---|
| | Genes (In at least one subsample) | | | | | | | |
| < 0.1 | 2066 (17.2%) | 513 (+4.3%) | 194 (+1.6%) | 59 (+0.5%) | -3 (-0.02%) | -38 (-0.3%) | 2791 (23.2%) | 12010 |
| [0.1, 0.5) | 1963 (37.4%) | 377 (+7.2%) | 69 (+1.3%) | 12 (+0.2%) | -9 (-0.2%) | -29 (-0.6%) | 2383 (45.4%) | 5249 |
| [0.5, 1) | 1150 (55.2%) | 111 (+5.0%) | 4 (+0.2%) | -14 (-0.6%) | -5 (-0.2%) | -17 (-0.8%) | 1229 (55.8%) | 2204 |
| [1, 10) | 2900 (63.2%) | 75 (+1.6%) | 6 (+0.1%) | -16 (-0.3%) | -14 (-0.3%) | -25 (-0.5%) | 2926 (63.8%) | 4586 |
| ≥ 10 | 975 (61.7%) | 46 (+2.9%) | 13 (+0.8%) | -1 (-0.06%) | 1 (+0.06%) | 1 (+0.06%) | 1035 (65.5%) | 1580 |
| | Genes (In 100 subsamples) | | | | | | | |
| < 0.1 | 766 (6.4%) | +444 (+3.7%) | +310 (+2.6%) | +232 (+1.9%) | +179 (+1.5%) | +144 (+1.2%) | 2075 (17.3%) | 12010 |
| [0.1, 0.5) | 653 (12.4%) | +487 (+9.3%) | +328 (+6.2%) | +208 (+4.0%) | +138 (+2.6%) | +134 (+2.6%) | 1948 (37.1%) | 5249 |
| [0.5, 1) | 510 (23.1%) | +255 (+11.6%) | +135 (+6.1%) | +88 (+4.0%) | +51 (+2.3%) | +39 (+1.8%) | 1078 (48.9%) | 2204 |
| [1, 10) | 1892 (41.3%) | +438 (+9.6%) | +190 (+4.1%) | +102 (+2.2%) | +79 (+1.7%) | +50 (+1.1%) | 2751 (60.0%) | 4586 |
| ≥ 10 | 780 (49.4%) | +106 (+6.7%) | +44 (+2.8%) | +39 (+2.5%) | +17 (+1.1%) | +20 (+1.3%) | 1006 (63.7%) | 1580 |
| | Junctions (In at least one subsample) | | | | | | | |
| < 0.1 | 6889 (10.9%) | +2454 (+3.9%) | +975 (+1.5%) | +277 (+0.4%) | -64 (-0.1%) | -182 (-0.3%) | 10349 (16.4%) | 63026 |
| [0.1, 0.5) | 8221 (11.9%) | 2959 (+4.3%) | 847 (+1.2%) | 240 (+0.3%) | -251 (-0.2%) | -320 (-0.5%) | 11794 (17.1%) | 68798 |
| [0.5, 1) | 5510 (13.3%) | 1196 (+3.1%) | 192 (+0.5%) | -119 (-0.3%) | -117 (-0.3%) | -149 (-0.4%) | 6153 (15.9%) | 38621 |
| [1, 10) | 13327 (13.2%) | 1071 (+1.1%) | -17 (-0.02%) | -218 (-0.2%) | -285 (-0.3%) | -177 (-0.2%) | 13701 (13.5%) | 101345 |
| ≥ 10 | 3918 (9.7%) | 450 (+1.1%) | 172 (+0.4%) | 44 (+0.1%) | -13 (-0.03%) | -5 (-0.01%) | 4566 (11.3%) | 40517 |
| | Junctions (In 100 subsamples) | | | | | | | |
| < 0.1 | 2061 (3.3%) | +1438 (+2.3%) | +1138 (+1.8%) | +937 (+1.5%) | +676 (+1.1%) | +679 (+1.1%) | 6930 (11.0%) | 63026 |

| | | | | | | | | |
|---|---|---|---|---|---|---|---|---|
| [0.1, 0.5) | 1934 (2.8%) | +1949 (+2.8%) | +1505 (+2.2%) | +1238 (+1.8%) | +871 (+1.3%) | +805 (+1.2%) | 8302 (12.1%) | 68798 |
| [0.5, 1) | 1605 (4.2%) | +1161 (+3.0%) | +750 (+1.9%) | +540 (+1.4%) | +415 (+1.1%) | +362 (+0.9%) | 4822 (12.5%) | 38621 |
| [1, 10) | 6301 (6.2%) | +2317 (+2.3%) | +1188 (+1.2%) | +882 (+0.9%) | +638 (+0.6%) | +560 (+0.6%) | 11886 (11.7%) | 101345 |
| ≥ 10 | 2491 (6.1%) | +447 (+1.1%) | +301 (+0.7%) | +263 (+0.6%) | +184 (+0.5%) | +197 (+0.5%) | 3883 (9.6%) | 40517 |

**Supplementary Table S6**. The increase of the mean number and the percentage of genes and junctions involved in alternative splicing with increasing sequencing depth in the Hypothalamus dataset. Column '50 M' indicates the mean number and the percentage of genes/junctions detected in the samples with a sequencing depth of less than 50 M reads. The column '200M' indicates the mean number and the percentage of genes/junctions detected in the samples with the highest sequencing depth of 300M reads. Grey shading indicates an increase of less than 1% of new detections.

| TPM | 50M | 50-100 | 100-150 | 150-200 | 200M | Total # of genes/ junctions |
|---|---|---|---|---|---|---|
| | Genes (In at least one subsample) | | | | | |
| < 0.1 | 1925 (18.7%) | 517 (+5.0%) | 250 (+2.4%) | 10 (+0.1%) | 2702 (26.2%) | 10303 |
| [0.1, 0.5) | 1509 (24.4%) | 454 (+7.3%) | 184 (+3.0%) | 20 (+0.3%) | 2167 (35.1%) | 6179 |
| [0.5, 1) | 1239 (43.9%) | 220 (+7.8%) | 82 (+2.9%) | 15 (+0.5%) | 1556 (55.1%) | 2825 |
| [1, 10) | 5929 (67.6%) | 431 (+4.9%) | 155 (+1.8%) | 28 (+0.3%) | 6543 (74.6%) | 8771 |
| ≥ 10 | 2531 (79.0%) | 85 (+2.7%) | 26 (+0.8%) | 26 (+0.8%) | 2668 (83.3%) | 3203 |
| | Genes (In 100 subsamples) | | | | | |
| < 0.1 | 1074 (10.4%) | +576 (+5.6%) | +383 (+3.7%) | +421 (+4.1%) | 2454 (23.8%) | 10303 |
| [0.1, 0.5) | 604 (9.8%) | +566 (+9.2%) | +388 (+6.3%) | +388 (+6.3%) | 1946 (31.5%) | 6179 |
| [0.5, 1) | 592 (21.0%) | +435 (+15.4%) | +242 (+8.6%) | +194 (+6.9%) | 1463 (51.8%) | 2825 |
| [1, 10) | 4222 (48.1%) | +1340 (+15.3%) | +501 (+5.7%) | +336 (+3.8%) | 6399 (80.0%) | 8771 |
| ≥ 10 | 2315 (72.3%) | +1544 (+6.4%) | +573 (+2.2%) | +395 (+1.8%) | 2650 (82.7%) | 3203 |

| | Junctions (In at least one subsample) | | | | | |
|---|---|---|---|---|---|---|
| < 0.1 | 8661 (15.6%) | +2939 (+5.3%) | +1275 (+2.3%) | +57 (+0.1%) | 12932 (23.4%) | 55356 |
| [0.1, 0.5) | 7869 (11.1%) | +3634 (+5.1%) | +1638 (+2.3%) | −19 (−0.03%) | 13122 (18.5%) | 70978 |
| [0.5, 1) | 7217 (15.9%) | +2392 (+5.3%) | +836 (+1.8%) | −36 (−0.08%) | 10409 (23.0%) | 45310 |
| [1, 10) | 38921 (21.8%) | +7237 (+4.1%) | +1840 (+1.0%) | +196 (+0.1%) | 48194 (27.0%) | 178510 |
| ≥ 10 | 15363 (23.0%) | 1795 (+2.7%) | 915 (+1.4%) | 466 (+0.7%) | 18539 (27.8%) | 66807 |
| | Junctions (In 100 subsamples) | | | | | |
| < 0.1 | 3642 (6.6%) | +2774 (+5.0%) | +2177 (+3.9%) | +2601 +(4.7%) | 11194 (20.2%) | 55356 |
| [0.1, 0.5) | 2177 (3.1%) | +2842 (+4.0%) | +2734 (+3.9%) | +3069 (+4.3%) | 10822 (15.2%) | 70978 |
| [0.5, 1) | 2221 (4.9%) | +2717 (+6.0%) | +2065 (+4.6%) | +2087 (+4.6%) | 9090 (20.1%) | 45310 |
| [1, 10) | 18321 (10.3%) | +11842 (+6.6%) | +7313 (+4.1%) | +6926 (+3.9%) | 44402 (24.9%) | 178510 |
| ≥ 10 | 10238 (5.4%) | +15422 (+5.4%) | +9408 (+3.1%) | +8755 (+2.7%) | 17742 (26.6%) | 66807 |

**Supplementary Table S7.** The results of NEASE with KEGG pathways for the deep-sequenced data sets

| **KEGG Pathway** | **P-adjusted** |
|---|---|
| **Adipose (pre-treatment)** | |
| Citrate cycle (TCA cycle) | 0.0052 |
| Propanoate metabolism | 0.0052 |
| Apoptosis | 0.0059 |
| PI3K-Akt signaling pathway | 0.0061 |
| p53 signaling pathway | 0.0078 |
| Small cell lung cancer | 0.0087 |
| Gastric cancer | 0.0107 |
| Pyruvate metabolism | 0.011 |

| | |
|---|---|
| Ferroptosis | 0.014 |
| Ras signaling pathway | 0.016 |
| Antigen processing and presentation | 0.018 |
| Prostate cancer | 0.02 |
| VEGF signaling pathway | 0.021 |
| Hepatocellular carcinoma | 0.025 |
| Vasopressin-regulated water reabsorption | 0.03 |
| Thyroid hormone signaling pathway | 0.038 |
| Non-small cell lung cancer | 0.04 |
| Valine, leucine and isoleucine degradation | 0.042 |
| Hepatitis B | 0.042 |
| Fatty acid degradation | 0.043 |
| Acute myeloid leukemia | 0.044 |
| **Adipose (post-treatment)** | |
| SNARE interactions in vesicular transport | 0.0056 |
| NOD-like receptor signaling pathway | 0.0059 |
| Herpes simplex infection | 0.0103 |
| Epstein-Barr virus infection | 0.0103 |
| TNF signaling pathway | 0.0103 |
| Toll-like receptor signaling pathway | 0.021 |
| Measles | 0.024 |
| Antigen processing and presentation | 0.026 |
| Endocytosis | 0.028 |
| mTOR signaling pathway | 0.031 |
| Proteasome | 0.031 |
| **Hypothalamus** | |
| Prion diseases | 0.0048 |

| | |
|---|---|
| Phagosome | 0.0055 |
| Gap junction | 0.007 |
| Oxidative phosphorylation | 0.0092 |
| Adherens junction | 0.023 |
| Glutamatergic synapse | 0.023 |
| Insulin secretion | 0.024 |
| Propanoate metabolism | 0.025 |
| Fatty acid degradation | 0.025 |
| Fructose and mannose metabolism | 0.025 |
| Leishmaniasis | 0.026 |
| Pancreatic secretion | 0.026 |
| Glycolysis / Gluconeogenesis | 0.026 |
| Hippo signaling pathway | 0.026 |
| Valine, leucine and isoleucine degradation | 0.026 |
| Protein processing in endoplasmic reticulum | 0.027 |
| Oocyte meiosis | 0.027 |
| Collecting duct acid secretion | 0.032 |
| Carbohydrate digestion and absorption | 0.044 |
| cAMP signaling pathway - | 0.046 |
| Human T-cell leukemia virus 1 infection | 0.048 |
